# A radial map of the budding yeast genome reveals novel organizational principles

**DOI:** 10.64898/2026.05.06.722996

**Authors:** Gertjan Laenen, Wing Hin Yip, Manuela Baquero Pérez, Axel Cournac, Magda Bienko, Angela Taddei

## Abstract

The eukaryotic genome is non-randomly organized within the nucleus, with positioning linked to function. Still, genome-wide radial maps are missing for the majority of experimental model systems. We adapted Genomic loci Positioning by Sequencing (GPSeq) to *Saccharomyces cerevisiae*, enabling high-resolution mapping along the nuclear center–periphery axis. GPSeq confirms known spatial features and shows that peripheral telomeres and centromeres impose long-range constraints extending up to 200 kb, restricting short chromosome arms from the nuclear interior. Telomere repositioning to the nuclear center, either artificially or during quiescence, reorganizes much of the genome through inward movement of sub telomeric regions and compensatory shifts of mid-arm chromatin outward. In quiescence, reduced centromere peripheral localization further alters genome organization. While transcription has a modest impact on radial positioning in all studied conditions, we uncover that in the absence of centromere or telomere constraints, GC-content functionally organizes chromatin in the nucleus.

**Graphical abstract:** The budding yeast genome is spatially organized in a manner highly dependent on the positioning of centromeres (*CENs*) and telomeres (*TELs*). Anchoring of these chromosome landmarks constrains the positioning of adjacent chromatin up to 200 kb within the same radial zone. Beyond this range, genome organization is non-random, with processes like transcription and features such as GC- content associated with specific radial positions in the nucleus.

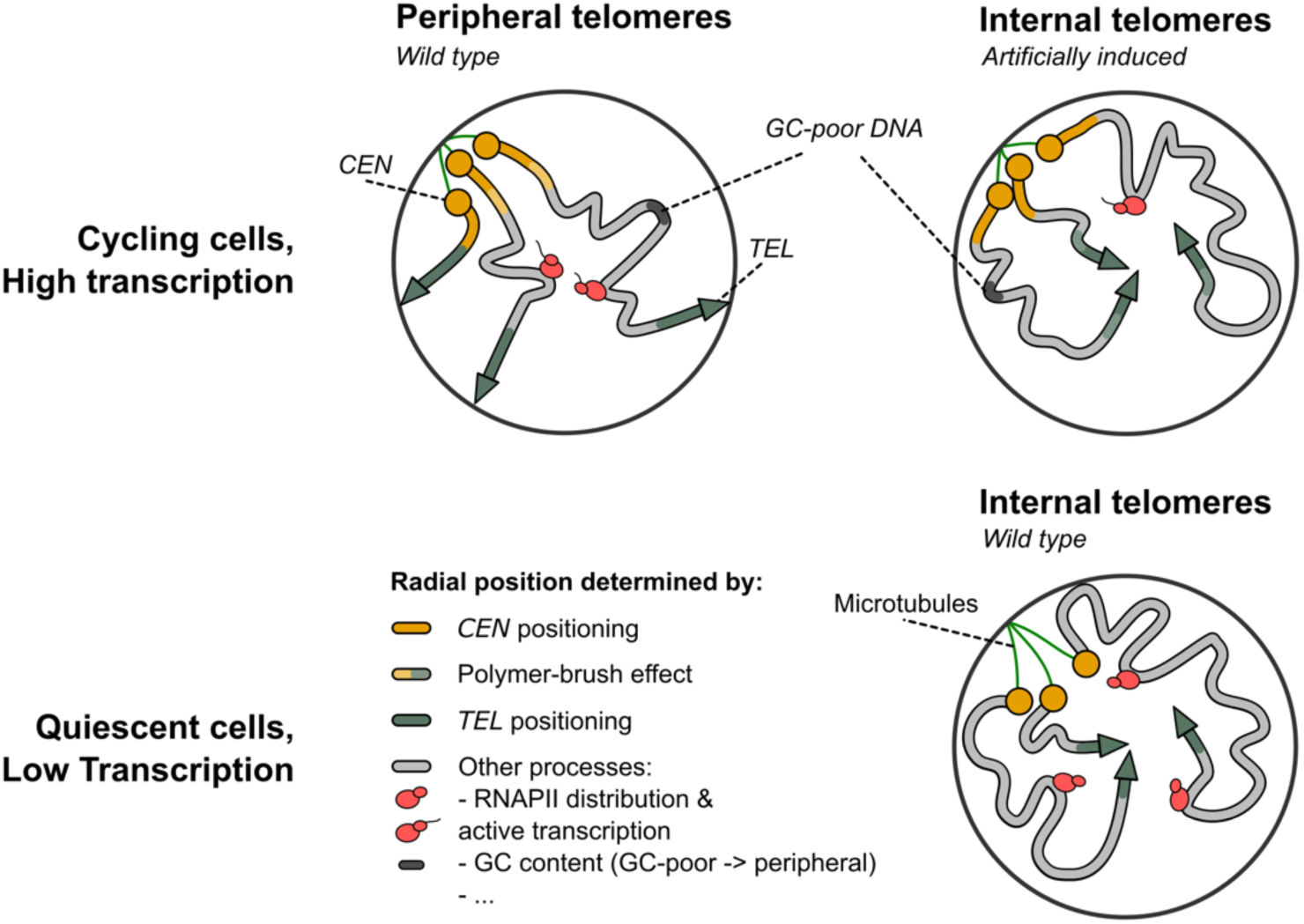

## Introduction

The eukaryotic genome is spatially organized in the nucleus in a non-random fashion. Large portions of the genome are physically anchored to the nuclear periphery, and nuclear processes ranging from transcription to DNA replication or genome maintenance either influence or are influenced by 3D genome architecture^1,2^. Nuclear processes shape the intranuclear arrangement of chromatin, yielding various functional subcompartments. Such subcompartments are marked by colocalization of specific DNA loci and, often, can be characterized by a preferential position relative to the nuclear periphery/center. One such subcompartment conserved in different cell types and across species is the nuclear rim or periphery, known to be enriched in DNA loci undergoing transcriptional silencing^3–5^. This region is contrasted by transcriptionally active loci, which have a tendency to occupy the nuclear interior^6,7^. Accordingly, when active genes are artificially tethered to the nuclear periphery, they tend to undergo downregulation^8–10^. *Vice versa,* genes often move toward the nuclear interior when de-repressed^11–14^. Although these general principles are well conserved from yeast to human cells, they don’t apply to all cell types or growth conditions^15,16^.

Genome architecture of the budding yeast *Saccharomyces cerevisiae* has been extensively studied in cycling cells and presents a well-defined organization. Throughout the cell cycle, centromeres remain attached to the spindle pole body (SPB), itself embedded within the nuclear envelope, which in turn restricts their movement into the nuclear interior^17–19^. At the other nuclear pole, opposite from the centromeres, *rDNA* repeats localize to the nuclear periphery, surrounded by a nucleolus that encompasses a third of the nucleus^20,21^. Silent chromatin in the budding yeast nucleus is localized at two cryptic mating type loci and at each of the 32 telomeres, which coalesce into 4-8 clusters, forming a subnuclear compartment that favors and restricts gene silencing to the nuclear periphery^1,3^. This organization is often referred to as ‘Rabl-like’, with telomeres positioned away from the centromeric cluster.

Importantly, the organization of the budding yeast genome is dynamic and undergoes changes according to the cellular environment. For example, in alternative growth conditions, a handful of genes has been shown to move toward the nuclear periphery alongside their activation^22–26^. In quiescent yeast, silent chromatin relocalizes inwards, with telomeres coalescing into a single hypercluster in the nuclear interior^16,27^. This dramatic reorganization is reminiscent of the internalization of heterochromatin in rod the cells of nocturnal animals^15^. Notably, during budding yeast quiescence, the majority of the genome is transcribed at very low levels^27–30^, but the relationship of this transcriptional change with radial positioning remains poorly understood, including for the few quiescence-specific active genes^27^.

Improvements in fluorescence microscopy and chromosome conformation capture techniques^16,31,32^, helped to shed light onto various aspects of nuclear organization in the yeast quiescent state. However, genome-wide large-scale reorganizations, characterized by the formation of the telomere hypercluster in the center of the nucleus, remain unexplored. Even during exponential growth on glucose—the most studied condition—the behavior of chromatin relative to the nuclear periphery/center is still elusive at genome-wide scale. While coarse maps of chromosomes XII^33^ and II^34^ have been generated by probing a relatively small number of engineered loci with fluorescence microscopy, strain construction and sequential imaging hinder high-resolution genome-wide coverage.

To address this knowledge gap, here we adapted Genomic loci Positioning by Sequencing (GPSeq)^35,36^ to map genomic loci along the center–periphery axis of the yeast nucleus at high resolution genome-wide. GPSeq exploits the gradual radial diffusion of restriction enzymes into the cell nucleus, by subjecting cells or nuclei to progressively increasing incubation times with one restriction enzyme. By amplifying and sequencing the genomic sequences surrounding the digested restriction sites, followed by comparison of the read coverage at each restriction site across all the digestion timepoints, a GPSeq score is generated for each genomic bin of a given length, effectively producing a genome-wide map of the radial placement of individual genomic bins^35,36^. We first validated our yeast GPSeq workflow by focusing on previously observed behaviors of the genome along the nuclear radius. We then characterized the typical radial placement of different regions of the yeast genome for which no prior knowledge has been available. We also applied GPSeq under conditions that drive telomeres inwards, revealing how radial changes across the whole genome accommodate the telomere hyper-clustering in the very center of the nucleus. In addition, we found that transcribed genes are slightly enriched at the nuclear center and found evidence supporting a context-dependent link between radial position and GC-content of DNA sequences.

## Results

### Genomic loci Positioning by Sequencing (GPSeq) in *Saccharomyces cerevisiae*

In order to generate GPSeq maps for the budding yeast genome, we first adapted the GPSeq protocol (previously applied to adherent human cells) to suspension budding yeast cells carrying a cell wall (**Fig. 1a**). The yeast GPSeq (yGPSeq) workflow relies on the progressive diffusion of a restriction enzyme through chromatin (here we used DpnII, with cutsites every ∼337 bp of the yeast genome), starting at the nuclear periphery all the way to the nuclear center. While mammalian nuclei typically have a radius of 5–10 μm, yeast nuclei often have a radius of ∼1 μm (**Supplementary Fig. 1a**), reducing the available nuclear volume by roughly 100–1000-fold. This drastic reduction in scale posed major challenges for adapting GPSeq to yeast, as enzyme diffusion takes place within a far smaller space. We assesed gradual enzyme diffusion and digestion by using the YFISH assay^35,36^ (**Fig. 1b**). In YFISH, dsDNA adaptors that carry a 5’-overhang are ligated to DpnII-digested cutsites. The other end of the adaptor is fork-like allowing for binding of fluorescent oligos to both ‘tines’. Through fluorescence microscopy, digested cutsites can then be visualized in the nucleus, allowing the assessment of the progression of enzyme digestion across different time points.

**Figure 1.**
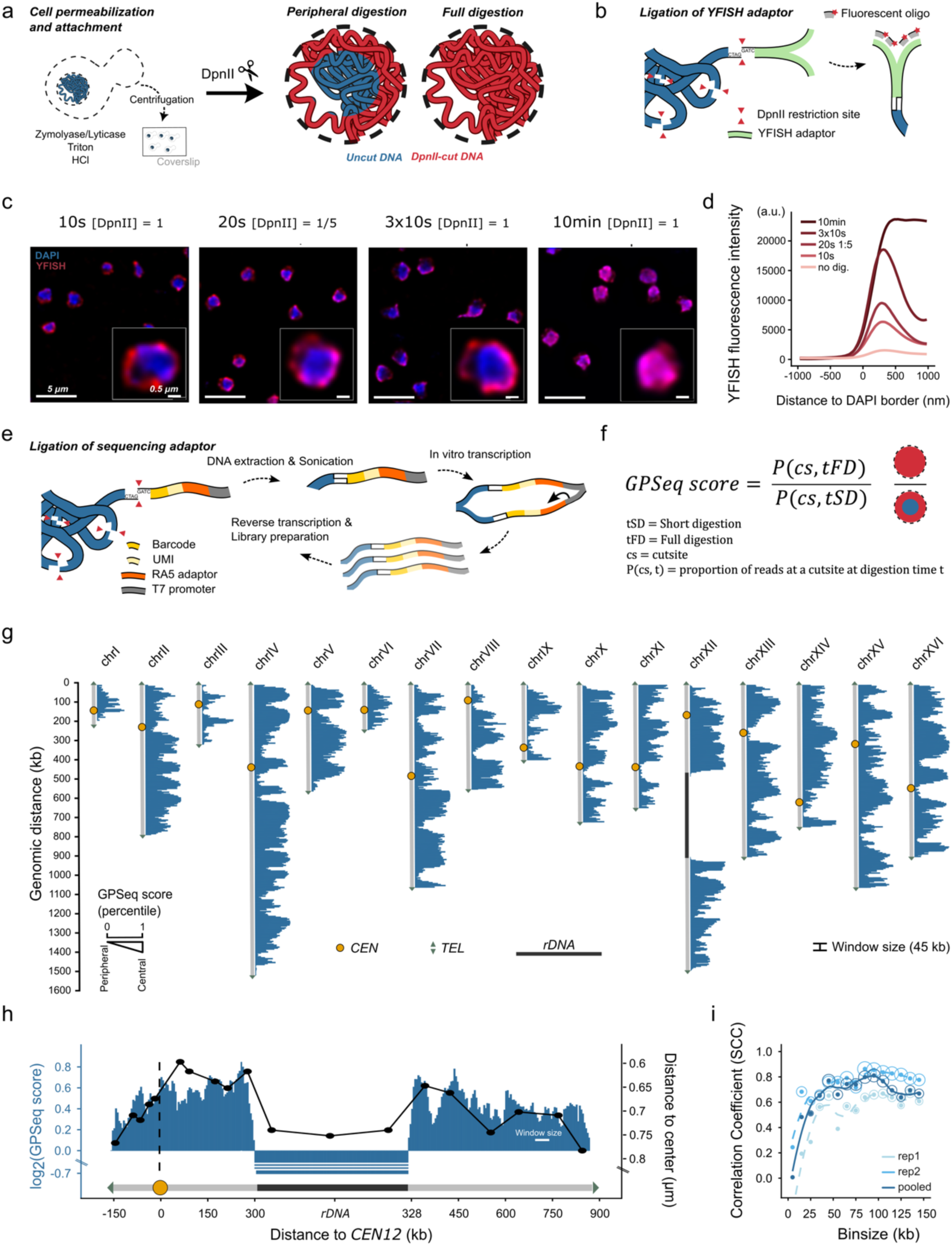
GPSeq in *Saccharomyces cerevisiae*. **a**: GPSeq workflow and visualization: Cells are permeabilized with cell wall digestion enzymes in suspension, then centrifuged onto slides. Afterwards, a mild detergent is used and HCl at low concentration, followed by *in situ* restriction digestion in one of two ways. **b**: Schematic illustration of post-digestion ligation of YFISH adaptors to the cutsites, allowing for the binding of fluorescent oligos. **c**: Images of cycling yeast cells, showing different short digestions and a long digestion. Deconvolution is used to enhance visibility of the peripheral signal (**see Methods**).**d**: Fluorescence intensity levels of YFISH signals from different digestion times and enzyme concentrations (10 min, 3×10s and 10s at the same concentration; 20s: at five-fold dilution; no digestion: 10 min without enzyme) in wild-type cycling cells. **e**: Schematic illustration of the ligation of the sequencing adaptor and sequential library preparation. **f**: GPSeq scores in budding yeast are calculated as the ratio of reads coming from the full digestion over the reads coming from the peripheral digestion, yielding values from low (peripheral) to high (central). **g**: Chromosome profile of GPSeq score percentiles for all chromosomes in the budding yeast genome, using 45 kb sliding windows (5 kb step). The sequence underlying *rDNA* repeats is artificially stretched for ∼400 kb to illustrate a more realistic occupancy of the *rDNA* repeats. **h**: Chromosome profile of *chrXII*, combining log_2_(GPSeq scores) with the point-median data of Albert *et al.* (2013) superimposed. The *rDNA* locus was repeated 30 times to reflect a more realistic genomic occupancy. GPSeq score is plotted in 45 kb windows, sliding in 5 kb steps. **i**: SCC between GPSeq scores and loci probed through FROS-imaging for *chrXII*, for increasing binsizes. Size of circles surrounding each point represented the −log_10_(p-value) associated with the correlation, only when p.value < 0.05.

We first tested various strategies to ensure progressive radial digestion of chromatin in the small nucleus of yeast, either by varying incubation time or enzyme concentration. A 10-minute digestion yielded a homogenously digested genome all across the nucleus resulting in a strong overlap between the YFISH and DAPI signals (**Supplementary Fig. 1b**). Next, we tried several approaches to limit the digestion to the nuclear periphery (**Fig. 1c-d and Supplementary Fig. 1c**). A single short burst of DpnII exposure (10s) resulted in a faint digestion signal restricted to the periphery of the DAPI signal. Diluting the enzyme (1:5) and exposing the cells to a slightly longer pulse of digestion (20s) resulted in a slightly stronger signal. Finally, repeating the 10s pulse of digestion three separate times yielded the brightest signal, producing a three-fold increase in fluorescence intensity relative to a single 10s pulse, while remaining highly restricted to the very periphery (**Fig. 1d**). While multiple approaches allowed for an enrichment of cuts at the nuclear periphery (**Fig. 1c-d and Supplementary Fig. 1c**), an intermediate digestion was difficult to visually assess. Hence, for GPSeq in *S. cerevisiae*, we decided to rely exclusively on a peripheral and a full digestion time point, which given the bulk nature of the assay yield a continuous range of the radial placement of DNA loci. In line with this approach, we previously found that relying on two most extreme time points of digestion in human cells already yielded accurate maps of radiality (unpublished data).

Having identified optimal digestion conditions for a peripherally restricted versus full digestion, we then substituted YFISH-compatible adaptors with adaptors allowing for *in vitro* transcription and sequencing of the genomic region surrounding the restricted sites (**Fig. 1e**). Following sequencing of yGPSeq libraries on Nextseq 2000, we performed mapping of the reads to specific DpnII cutsites along the genome (**see Methods**). Between all three types of peripheral digestions (10”, 20” at 1:5 enzyme dilution, and 3 x 10”), the reads mapped to individual DpnII cutsites correlated strongly with one another (Spearman’s correlation coefficient, SCC: 0.90-0.97; **Supplementary Fig. 1d**). Therefore, we treated these as technical replicates and pooled the read counts from these three conditions together before calculating the GPSeq scores. GPSeq scores were calculated in a manner similar to what we previous described^36^, using the two digestion time points as reference and dividing the number of reads obtained from the fully digested sample by the number of reads from the peripherally digested sample (**Fig. 1f and Methods**). This yielded values of the GPSeq score, ranging from low (peripheral loci) to high (central loci) (**Fig. 1g**). To assess the GPSeq score robustness, we compared biological replicates at different bin sizes (5-150 kilobases, kb) and found that the GPSeq scores consistently correlated (SCC: ∼0.75-0.8) across biological replicates (**Supplementary Fig. 1e**). For most visualizations and quantifications in this work, we used genomic bins in the range of 20–80 kb, unless noted otherwise. As for human chromosomes, we observed that the GPSeq score in yeast also fluctuates along individual chromosomes, indicating local differences in radial positioning (**Fig. 1g**).

To benchmark GPSeq scores and the local differences we observe, we compared the GPSeq profiles to microscopy-based radial maps previously described for a subset of fluorescently tagged loci along the *S. cerevisiae* chromosome XII^33^ (**Fig. 1h**). The two datasets correlated strongly (SCC: 0.8 at 45 kb sliding windows), with GPSeq reproducing key features of the radial distribution of the yeast *chrXII*, such as a relatively central position of its centromere, local arm-level fluctuations and the peripheral placement of its *rDNA* repeats (**Fig. 1h and Supplementary Fig. 1f**). The relatively central position of its centromere potentially reflects perinuclear tethering of the rDNA at the pole opposite to the centromere cluster^20^, effectively pulling *CEN12* toward the nuclear interior. Noteworthy, the correlation between these two datasets is statistically significant at window sizes of 40 kb and above, peaking at around 80 kb (**Fig. 1i**). A similar though weaker trend is observed when comparing GPSeq scores with microscopy-based measurements of locus association with the nuclear periphery by Dultz *et al.* (2016)^34^ (**Supplementary Fig. 1g-h**). Altogether, we show how minor changes to the GPSeq workflow allow for the generation of a high-resolution genome-wide map of radial localizations in budding yeast.

### Peripheral centromeres and telomeres radially organize large portions of the genome

We found centromeres and telomeres to be relatively peripheral (**Fig. 1g and Fig. 2a**), in line with previous studies^1,3,19^. To ensure adequate coverage of repetitive subtelomeric regions, we allowed multimapping reads within 13 kb of chromosome ends. This approach substantially increased the amount of available cutsites and improved the signal at telomeres without affecting the more internal loci (**Supplementary Fig. 2a-b**). Although both centromeres and telomeres are significantly more peripheral than the average locus, centromeres show a milder tendency in comparison to telomeres. These results are consistent with fluorescence microscopy studies showing that centromeres are indirectly connected to the spindle pole body through microtubules, placing them 200–300 nm away from the nuclear envelope^19^. Telomeres, by contrast, are anchored more directly to the nuclear envelope^1^.

**Figure 2.**
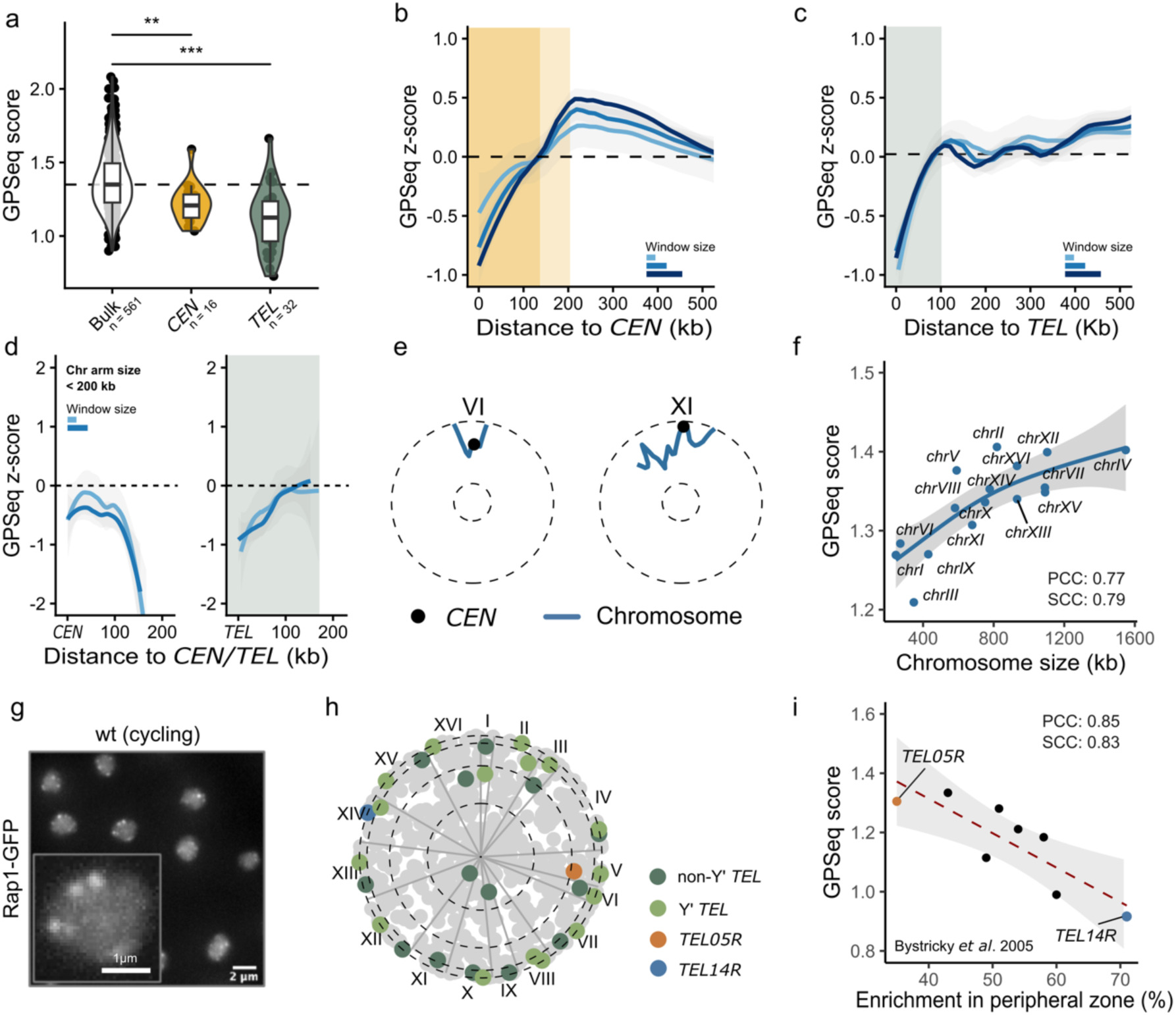
The Rabl-like organization in *S. cerevisiae* imposes long range radial constraints on chromosome arms. **a**: GPSeq scores of telomeres and centromeres compared to the bulk of the yeast genome (binsize: 20 kb). The dashed line represents the genome-wide median GPSeq score. Significance is shown for the results of a Wilcoxon rank-sum test: **: p-value < 0.01, ***: p-value < 0.001. **b**: Average GPSeq z-scores extending from the centromere. Line color in represents different window sizes (scales relative to x-axis). Smallest binsize shows non-overlapping (20 kb) windows, while the other two represent overlapping sliding windows, respectively 45 kb and 80 kb, both with a 5 kb step. Shaded sections of the graph represent different sections of increasing GPSeq scores. **c**: Same as in **b** but now showing average GPSeq z-scores extending from the telomere. **d**: Same as in **b-c**, but for chromosome arms smaller than 200 kb in length. **e**: Radial chromosome profile plots showing GPSeq scores binned at 75 kb windows sliding in steps of 25 kb. **f**: Scatter plot showing average GPSeq scores per chromosome relative to their size. Pearson Correlation Coefficient (PCC) and Spearman Correlation Coefficient (SCC) show significant correlations: p-value < 0.05. **g**: Fluorescence microscopy image of the telomere binding Rap1-GFP protein in cells growing exponentially on glucose (wt). Scale bars represent 1 and 2 μm respectively. **h**: Pizza plot highlighting telomeric-most 20 kb for telomeres either containing (Y’ *TEL*) or not containing (non-Y’ *TEL*) a repetitive Y’ element. Bins underlying telomeric regions probed in Bystricky *et al.* (2005) are highlighted for *TEL05R* and *TEL14R*, binsize = 20 kb non-overlapping bins. **i**: Scatter plot, showing the correlation between GPSeq scores and peripheral enrichment of telomeric loci probed through FROS (Fluorescent repressor-operator system) in Bystricky *et al.* (2005). PCC and SCC are significant, p-value < 0.02.

We noticed that GPSeq scores increased with distance from centromeres and telomeres, rising from peripheral values toward the genome-wide average within the first 100–150 kb (**Fig. 2b-c**). While telomere-adjacent GPSeq scores plateaued at the genome-wide average, centromere-adjacent scores kept increasing, surpassing the genome-wide average at 100 kb, peaking at ∼200 kb from the centromeres (**Fig. 2b**). Of note, genomic regions up to this distance from the centromeres show enhanced proximity in Hi-C (**Supplementary Fig. 2c-d**), an observation previously attributed to a polymer-brush effect emanating from the centromere^37^. We thus conclude that, on average, the brush effect projects 200 kb of centromere proximal chromatin of the 32 chromosome arms toward the nuclear interior. For long chromosome arms, this effect slowly dissipates for genomic loci located between 200 and 500 kb from centromeres.

Chromosome arms smaller than 200 kb (**Fig. 2d**) transition directly from the radial position of the telomere to the position of the centromere, with minimal deviation toward the nuclear center. Hence globally, small chromosomes and small chromosome arms are retained at the nuclear periphery (**Fig. 2e**). This behavior scales with chromosome size as observed from the correlation plot between chromosome size and GPSeq score (**Fig. 2f**), where mainly the largest chromosomes carry bins that are characterized by high GPSeq scores (central). This is consistent with the physical properties of the chromatin fiber, inferred from live imaging and HiC data^19,38–40^. For example, in the W303 yeast background^41^, chromosome I is ∼246 kb long, with its centromere positioned 152 kb from one end. Assuming both centromere and telomeres are peripherally anchored, the longest arm (*chrIL*) can extend at most ∼76 kb toward the nuclear interior. Using a worm-like chain model of chromatin (R²(S) = 2PS[1 − P/S(1 − e^(−S/P))]) with persistence length P (52–85 nm^40^) and contour length S calculated from genomic distance using a chromatin compaction of 53–65 bp/nm^40^, we estimate that the longer arm can reach ∼341–479nm, toward the center of the nucleus. Thus, given a nuclear radius of ∼1 μm, neither arm can fully extend to the nuclear center.

We then further explored potential differences between individual centromeres and telomeres. Fluorescence microscopy of the telomeric factor Rap1 shows strong clustering and localization at the nuclear periphery, albeit there is variation in cluster number and localization between cells (**Fig. 2g**). While microscopy captures this cell-to-cell heterogeneity aspect of telomere positioning being blind to the identity of individual telomeres (in the context of typical telomere stainings), we used our GPSeq data to reveal the typical radial placement of every single telomere. We plotted telomeric bins across all chromosomes in a “pizza”-like radial representation, where each chromosome is represented by a sector of the radial plot (**Fig. 2h**). Most telomeric bins showed low GPSeq scores, consistent with their peripheral localization, however some deviated from this pattern (such as *TEL09R* and *TEL11L*). Interestingly, in the W303 genome, 18/32 subtelomeres contain the repetitive Y’ element, and we observed that those telomeres show more peripheral GPSeq scores than telomeres without it (**Fig. 2h and Supplementary Fig. 2e**). We excluded a possible bias stemming from cutsite-availability, as there is no notable difference between Y’ element-bearing and -lacking telomeres for how many DpnII cutsites they carry (**Supplementary Fig. 2b**). Importantly, for a subset of telomeres whose radial localization was previously quantified by microscopy in the W303 background^42^, those measurements correlated highly with subtelomeric GPSeq scores (SCC = –0.83; **Fig. 2i**). These results confirm that GPSeq captures both a general peripheral tendency and the heterogeneity of telomere positioning.

In conclusion, yGPSeq expands our understanding of telomere and centromere positioning in the yeast nucleus. Our data show that these peripheral landmarks constrain the chromosome-arm over distances of up to 200 kb. In addition, the genome-wide resolution of yGPSeq reveals substantial heterogeneity in the spatial behaviour of these landmarks, extending previous observations to the whole genome.

### The internalization of telomeres affects genome organization through direct and compensatory effects

We next examined how radial genome architecture changes when telomere localization differs from the canonical Rabl-like configuration as observed in quiescent cells^16^. Quiescent yeast cells were sorted from a seven-day stationary phase culture as previously described^43^. This metabolic state represents an inactive yet stress-resistant phase associated with enhanced longevity^43^. Quiescent cells display smaller nuclei, very sparse transcription ^27,28,30^ and, as we reported previously, their telomeres coalesce into a single hypercluster near the nuclear center^16^ (**Fig. 3a**). To control for direct effects coming from transcriptional downregulation in quiescence, which might affect radial organization in addition to the telomere hyperclustering, we also profiled cells overexpressing the silencing Sir3-A2Q mutant protein while cycling. Sir3-A2Q overexpression drives telomere clustering into the center of the nucleus (**Fig. 3a**) without extending heterochromatin domains or leading to genome-wide differences in transcription, except for few subtelomeric genes^44–46^.

**Figure 3:**
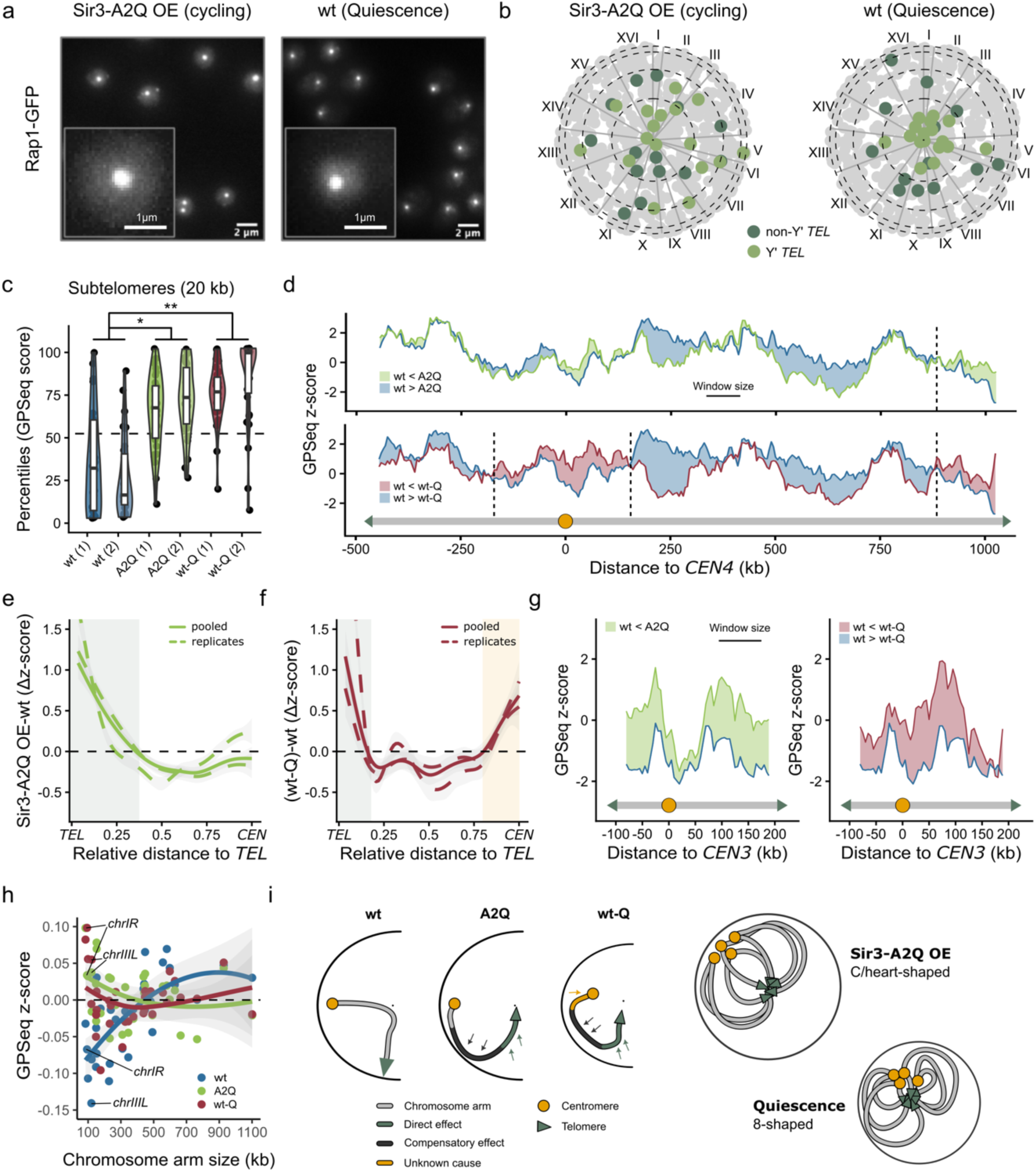
Telomere internalization affects genome organization through direct and compensatory effects. **a**: Fluorescence microscopy of the telomere binding Rap1-GFP protein in Sir3-A2Q OE and quiescent cells. **b**: Pizza plot highlighting telomeric-most 20 kb for telomeres either containing (Y’ *TEL*) or not containing (non-Y’ *TEL*) a repetitive Y’ element for A2Q and wt-Q. Binsize = 20 kb non-overlapping bins. **c**: GPSeq score percentiles for subtelomeric reads (20 kb bins) showing replicates for each condition. Pairwise significance was assessed from estimated marginal means derived from a generalized linear mixed-effects model, with biological replicate included as a random effect: *: p-value < 0.05, **: p-value < 0.01.**d:** Chromosome profile for *chrIV*, showing GPSeq z-scores for wt and A2Q/wt-Q. Difference between z-scores is colored. Windows of 80 kb were used, sliding in steps of 5 kb. **e:** Average difference in GPSeq z-score between A2Q and wt visualized over length normalized chromosome arms. Shaded sections of the graph represent different sections of the arm that are centralized on average.**f:** Average difference in GPSeq z-scores between wt-Q and wt visualized over length normalized chromosome arms. Shaded sections of the graph represent different sections of the arm that are centralized on average.**g:** Chromosome profile plots for *chrIII*. GPSeq z-scores for wt and A2Q/wt-Q. Difference between z-scores is colored. **h:** Average z-scores for different chromosome arm sizes. Different conditions are colored (wt:blue, A2Q:green, wt-Q:red).**i:** Model of genome reorganization in A2Q (C/heart-shaped, top) and wt-Q (8-shape, bottom).

We applied our newly adapted GPSeq protocol to both cycling Sir3-A2Q overexpressing cells (A2Q) as well as quiescent cells (wt-Q) (**Supplementary Fig. 3a-c**). While the cycling A2Q nucleus is of a similar size to cycling wild-type nuclei (wt), the wt-Q nucleus is significantly smaller (**Supplementary Fig. 3d**), resulting in a more difficult control of the DpnII enzyme diffusion. Despite this, we could obtain two distinct staining patterns: a peripheral YFISH staining upon a short pulse of digestion, as well as a pan-nuclear YFISH staining, even in the wt-Q cells (**Supplementary Fig. 3a-c**). In line with this, we obtained sequencing libraries that revealed reproducible GPSeq scores from both cell types (SCC: 0.7-0.8) at bin sizes of 5-150 kb (**Supplementary Fig. 3e**). Comparing GPSeq scores between all three conditions (wt, A2Q, wt-Q), we could observe a clustering of those from wt and A2Q, and a separate cluster for wt-Q cells (**Supplementary Fig. 3f**), indicating globally similar radial maps between wt and A2Q genomes, and a distinct radial organization genome-wide in quiescence.

Consistent with previous reports and with our fluorescent microscopy data (**Fig. 3a**), in both the A2Q and the wt-Q cells, GPSeq mapped telomeres to be central (**Fig. 3b**). Quantification across biological replicates showed a clear difference relative to wt, with GPSeq scores of telomeres being significantly more central in both A2Q and wt-Q in comparison to wt (**Fig. 3c**). To further explore how telomere repositioning influences chromosomal architecture, we examined GPSeq z-score profiles (**see Methods**) along an individual chromosome (**Fig. 3d**). Averaging differences of z-scores between wt and the two conditions for all chromosome arms, we find that approximately 150 kb windows adjacent to telomeres are more centrally localized in A2Q (**Supplementary Fig. 3g-h**), while in wt-Q this difference spans 50-100 kb (**Supplementary Fig. 3i-j)**. The difference in nuclear size could explain this difference, as the smaller wt-Q nucleus would physically require less chromatin to stretch from the nuclear center to the nuclear periphery.

In addition to telomeres pulling adjacent parts of chromosome arms toward the nuclear center, in both conditions, this repositioning of the telomeres is accompanied by a seemingly compensatory decrease in GPSeq scores over the mid-arm regions, indicating that chromatin in these regions becomes more peripheral than in wt (**Fig. 3d-f and Supplementary Fig. 3g-k**). This pattern could reflect a structural constraint: as telomeres move inwards, chromosome arms might be pushed outwards to accommodate the new configuration. While this feature is shared between A2Q and wt-Q, a key difference emerges around the centromeres. In wt-Q, the centromeres exhibit significantly higher (more central) GPSeq z-scores compared to wt, whereas in A2Q they remain relatively unchanged (**Fig. 3e-f and Supplementary Fig. 3k-n**). The release of centromeres from the nuclear periphery impacts the adjacent chromatin up to ∼125 kb on each side of the centromere (**Supplementary Fig. 3k**). Of note, while telomeres move into the nuclear center in wt-Q cells, centromeres lose peripheral location, but do not become more central than the average locus (**Supplementary Fig. 3l-m**).

For small chromosomes such as *chrI* and *chrIII*, whose arm lengths are < 250 kb, telomere internalization results in nearly the entire chromosome acquiring higher GPSeq z-scores in A2Q and wt-Q cells in comparison to wt (**Fig. 3g**). In wt-Q cells, the concurrent loss of the peripheral location of centromeres further accentuates this effect, leading to the disproportionate centralization of small chromosomes. To estimate the relationship between chromosome size and nuclear position, we compared mean GPSeq z-scores per chromosome arm to the length of the arm (**Fig. 3h**). In wt cells, arm length correlates positively with GPSeq z-scores (SCC: 0.70), reflecting the potential to occupy a more central position for large chromosome arms as discussed for chromosomes earlier (**Fig. 2d**). This relationship is lost or even inverted in A2Q (SCC: –0.34) and wt-Q (SCC: –0.14), consistent with the centralization of small chromosome arms and a compensatory peripheral displacement of internal segments of the bigger arms. The slightly stronger inversion in A2Q may stem from the persistence of peripheral centromeres in this condition, enforcing stronger bending of long chromosome arms.

These observations support a model of genome re-organization accompanying telomere repositioning where the radial placement of centromeres affects the type of reorganization (**Fig. 3i**). In the A2Q cells, telomeres move toward the nuclear interior while centromeres remain at the periphery, forcing chromosome arms to bend inwards and producing an overall “C/heart-shaped” nuclear organization. In the wt-Q cells, telomeres accumulate in the center, centromeres release from the periphery, while the center of the chromosome arms bends outwards potentially resulting in a more compact “8-shaped” configuration. Altogether, we observe how telomere reorganizations directly centralize roughly one-third of the genome in either condition (**Fig. 3e and Supplementary Fig. 3o-p**), with proportional peripheral-wards displacement of mid-arm chromatin compensating for this change.

### Active transcription is enriched toward the center of the budding yeast nucleus

We then wondered about the observed variation of the radial position of DNA loci along the arms of large chromosomes. We focused on the large chromosomes given that the short ones seem to be under a major influence of their anchor points and show less unrelated variability of the GPSeq scores along their arms.

We first studied transcription as a potential contributor to that variation, given that it has been shown to significantly interplay with the radial placement of genes in metazoans^6,7^. To this aim we have integrated our GPSeq data with various omics data previously generated for budding yeast (**see Methods**). Genome-wide correlations between GPSeq scores and RNAPII binding, transcription levels, or transcription-associated histone marks are weak (SCC: 0–0.2; **Supplementary Fig. 4a**). Nevertheless, we found features associated with active transcription to be consistently enriched in the central portion of the nucleus (**Fig. 4a**), in clear contrast to the peripheral accumulation of centromere- and telomere-associated factors (H3S10ph & Sir3). Here, we used Sir3 binding, which is mainly found at subtelomeres and silenced mating type loci, as a proxy for transcriptionally silenced genomic windows^1,3^. Interestingly, the central enrichment of features associated with active transcription persists after excluding subtelomeres from the analysis (**Supplementary Fig. 4b**) and in conditions where telomeres are internalized (A2Q, **Fig. 4b**), and even in the low-transcription state of quiescence cells (**Fig. 4c**).

**Figure 4:**
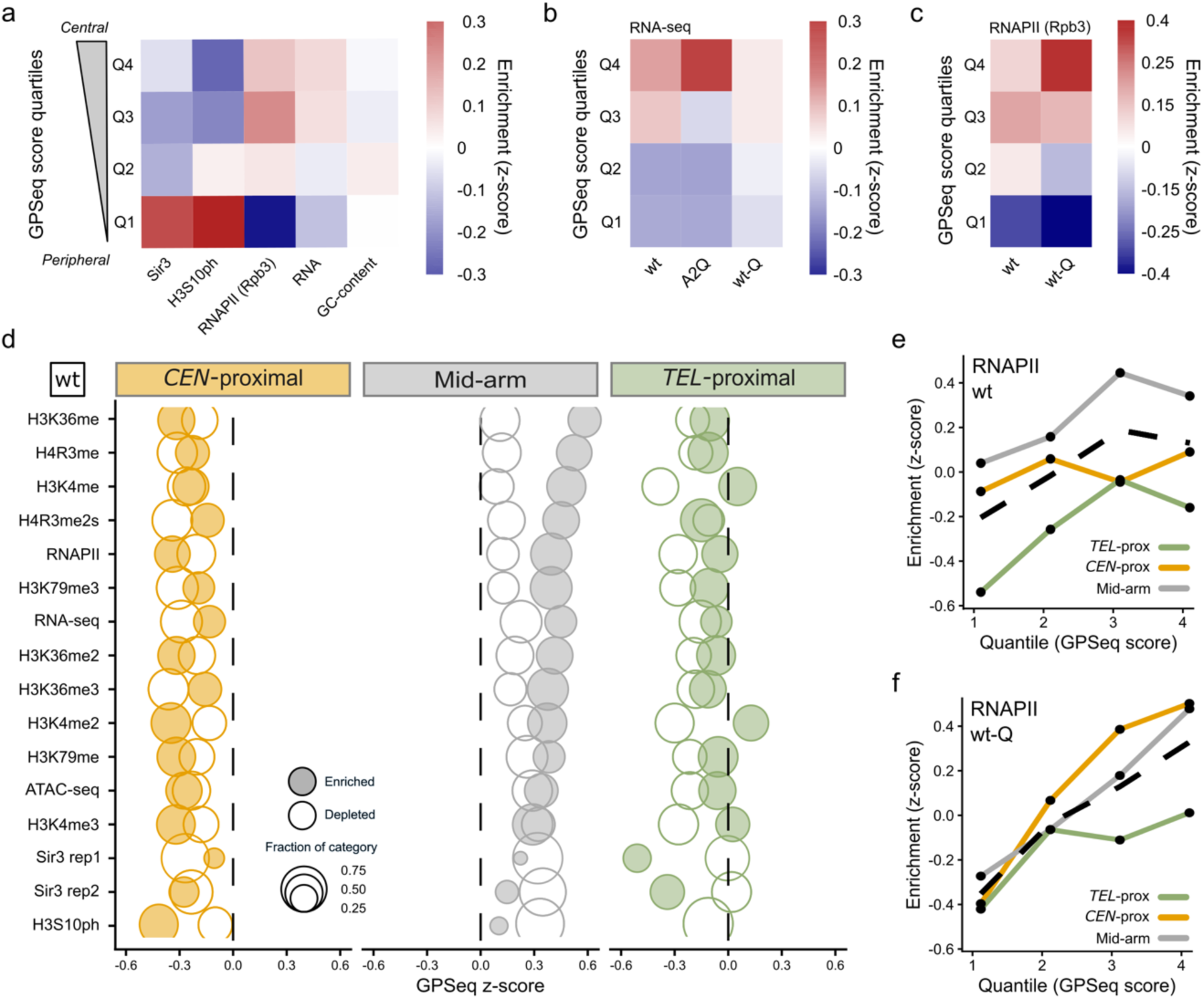
Active transcription is enriched toward the center of the budding yeast nucleus. **a**: Average feature z-score per GPSeq score quartile. Calculated from 45 kb windows sliding with 5 kb steps. The H3S10ph dataset was used from Weiner *et al.* (2015), RNA-seq data from Hocher *et al.* (2018) and RNAPII data from Baquero Pérez *et al.* (2025). **b**: Average z-score for RNA-seq counts per condition, shown for GPSeq score quartiles. Calculated from 45 kb windows sliding with 5 kb step. RNA-seq data for wt and A2Q was used from from Hocher *et al.* (2018) and RNA-seq data in wt-Q from Baquero Pérez *et al.* (2025). **c**: Average z-score for RNAPII (IP) counts per condition (wt, wt-Q), shown for GPSeq score quartiles. Calculated from 45 kb windows sliding with 5 kb step. RNAPII data was used from Baquero Pérez *et al.* (2025).**d**: Difference in median GPSeq z-score for regions enriched or depleted for a chromatin feature, features are ordered by GPSeq score distance between enriched and depleted for mid-arm chromatin. Histone modification data was used from Weiner *et al.* (2015), RNA-seq data for wt and A2Q from Hocher *et al.* (2018) and RNA-seq data in wt-Q and RNAPII data from Baquero Pérez *et al.* (2025). Shown for wt. Calculated from 45 kb sliding windows (5 kb step), bins are categorized by their proximity to *CEN* (634 bins)/ *TEL* (636 bins)/ mid-arm (1025 bins). **e**: Average RNAPII (IP) z-score over different GPSeq score quartiles for wt. Bins are separated by proximity to chromosome landmarks as in D. Dashed line represents the genome-wide average. RNAPII data was used from Baquero Pérez *et al.* (2025). **f**: Average RNAPII (IP) z-score over different GPSeq score quartiles for wt-Q. Bins are separated by proximity to chromosome landmarks as in D,E. Dashed line represents the genome-wide pattern. RNAPII data was used from Baquero Pérez *et al.* (2025).

We then dissected regional differences along individual chromosomes to identify regions where the link between radial location and gene expression is most pronounced. We compared centromere-proximal (*CEN* ±125 kb), telomere-proximal (*TEL* +150 kb), and mid-arm regions. In wt, active transcription (from RNA-seq and RNAPII binding) tends to be higher in more central GPSeq score quartiles for mid-arm and *TEL*-proximal regions, while less so for regions near centromeres (**Supplementary Fig. 4c**). A similar tendency was found in the A2Q cells (**Supplementary Fig. 4d**). Accordingly, when we intersected our radiality maps of wt cells with various epigenetic marks, we observed an interesting gradient of enrichment especially for the mid-arm level (but also to some extent for the *TEL*-proximal regions). Chromatin marked by H3K36me, H4R3me, H3K4me as well as H4R3me2s showed the strongest association with centrally located bins, followed by RNAPII and H3K79me3-associated bins. Of note, the ATAC-seq signal was not as strongly linked to the center of the nucleus, and finally moving toward the periphery we observed H3S10ph and Sir3 enrichment associated with low GPSeq scores, as expected (**Fig. 4d**).

Lastly, we focused on the interplay between radiality and transcription in the wt-Q cells, where genome expression is low and one-third of gene promoters are marked by intergenic accumulation of RNAPII^27^. While in quiescence, there was no enrichment/depletion of transcriptional activity in the consecutive quartiles of the nuclear radius (based on RNA-seq; **Fig. 4b and Supplementary Fig. 4e**), RNAPII was enriched in the central quartile, and depleted in the most peripheral one (**Fig. 4c**), and RNAPII binding showed a relatively strong correlation with GPSeq scores genome-wide (SCC: 0.34). This effect occurs for both the genic and intergenic binding of RNAPII (**Supplementary Fig. 4f**). Importantly, a major difference with cycling cells could be observed in *CEN*-proximal regions, where we observed a strong association of RNAPII binding with a more central positioning (**Fig. 4e-f**), possibly reflecting the lack of constraint imposed by centromeres in this condition.

### Inserting a GC-poor DNA is sufficient to shift a locus to the nuclear periphery and promote gene silencing

In the original application of GPSeq to human cells, chromosome size, gene density, gene expression and GC-content were probed as predictors of GPSeq scores, with GC-content revealing to be the strongest one^36^. Globally, the cycling wild-type budding yeast genome does not show any correlation between GC-content and GPSeq scores (**Fig. 5a**). However, in conditions where the telomeres are detached from the nuclear periphery (A2Q, wt-Q), correlations of GPSeq scores with transcription and GC-content increase significantly, lowering the effect imposed by proximity of chromosome landmarks (*CEN*, *TEL* or *rDNA*). In the A2Q cells specifically, GC-content seems to explain as much variation genome-wide, as is explained by landmark proximity in wt (**Fig. 5a**). Since both wt and A2Q are similar in their growth rate and gene expression, we decided to use their comparison to understand how relevant the effect of the GC-content might be on genome organization in the budding yeast nucleus.

**Figure 5:**
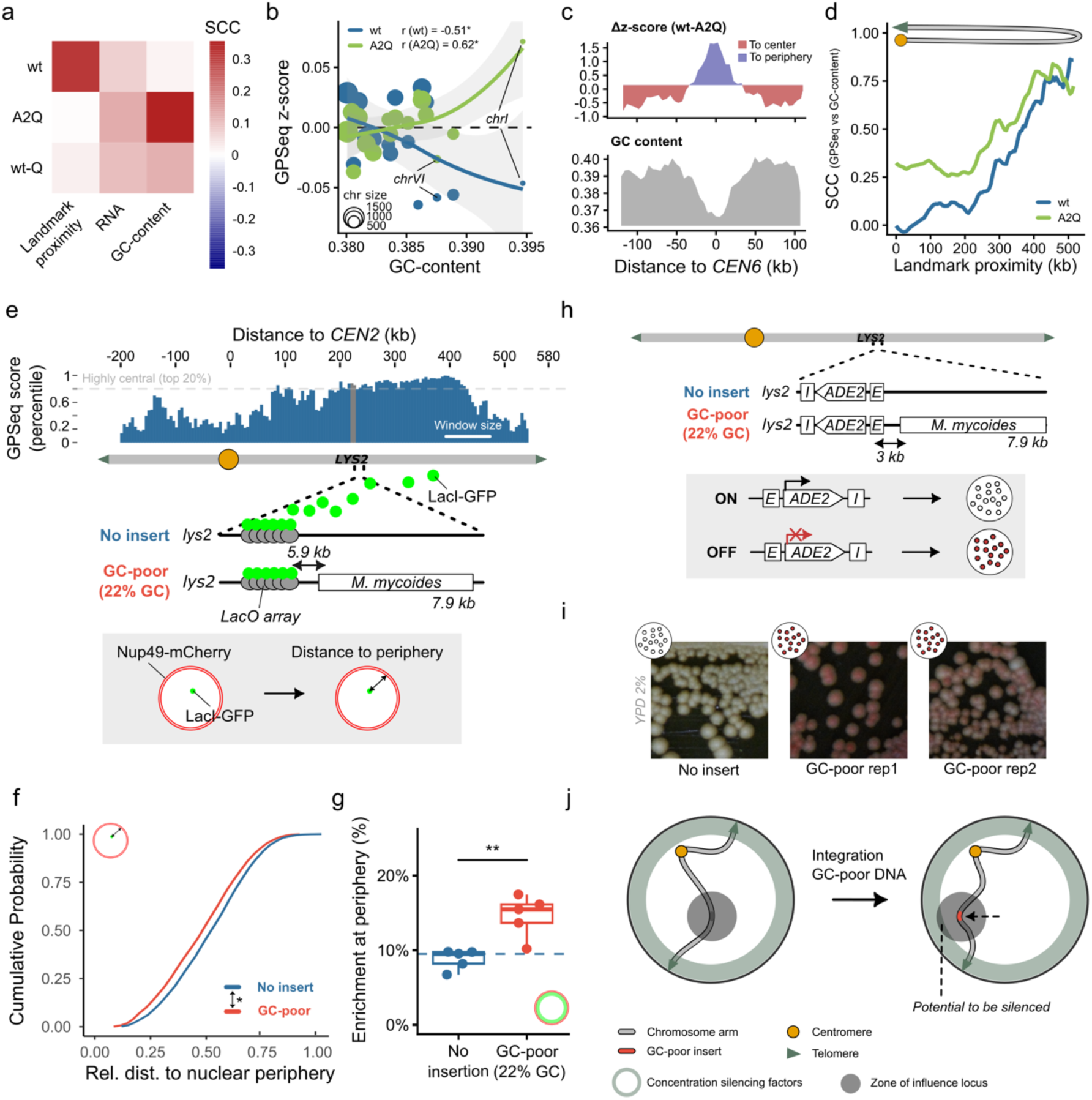
Inserting a GC-poor DNA is sufficient to shift a locus to the nuclear periphery, favoring gene silencing. **a**: SCCs between GPSeq score, landmark proximity, RNA-seq and GC-content. Calculated for 45 kb sliding windows (5 kb step). **b**: Scatterplot showing median GPSeq score (z-score) versus GC content for all chromosomes in A2Q and wt. Chromosome arm size is visualized as pointsize. **c**: Difference in z-score between GPSeq scores from wt and A2Q for *chrVI* shown above a graph showing GC-content over *chrVI*. Calculated for 45 kb sliding windows (5 kb step). **d**: Correlation between GC-content and GPSeq score shown while sequentially removing bins nearest to chromosome landmarks (*CEN*, *TEL* and *rDNA*). Correlation shown for 45 kb sliding windows (step: 5 kb). **e**: Chromosome profile of *chrII* (80 kb window, 5 kb step), showing the *LYS2* locus, with the lacO array insertion site used for its localization. **f**: Cumulative distributions of relative distance to the nuclear periphery for both the *lys2::lacO* (n=5) or *lys2::lacO M. mycoides* (7.9 kb) loci (n=5). Pairwise significance (Bonferroni post-hoc comparison) was assessed from estimated marginal means derived from a generalized linear mixed-effects model, with biological replicates included as a random effect: **: p-value < 0.01.**g**: Proportion of *lys2::lacO* (n=5) or *lys2::lacO M. mycoides* (7.9 kb, n=5) loci found at the periphery (n=5). The nuclear periphery is marked as the peripheral most third of the nuclear area. Significance is shown for the results of a Wilcoxon rank-sum test: **: p-value < 0.01.**h**: Scheme of *chrII* with *LYS2* locus, showing integration site of the silencing reporter and experimental setup of silencing assay. **i**: Ectopic silencing assay (ESA): colonies grown on YPD 2% agar plates at 30°C. Two replicates of GC-poor integration are shown in comparison to a control lacking the *M. mycoides* insert. **j**: Schematic representation of the potential mechanism: slight peripheralization allows the *lys2::lacO M. mycoides* (7.9 kb) locus to come into contact with silencing factors enriched at the nuclear periphery.

Among individual chromosomes, we observed a negative correlation between GC-content and GPSeq scores in wt (**Fig. 5b**), with small GC-rich chromosomes being retained at the nuclear periphery by telomere anchoring. Of note, this is the opposite to mammalian cells^36^. In A2Q, this trend inverts, with small GC-rich chromosomes being more central (**Fig. 5b**). Even more, the repositioning of individual sections of chromosomes in A2Q relative to wt follows GC-content (*chrVI*, **Fig. 5c**). These observations in A2Q suggest two potential sources for the correlation between GC-content and GPSeq scores. Either internalization of GC-rich *TEL*-proximal regions generates this increased correlation, or the peripheral anchoring of centromeres and telomeres in wt masks an intrinsic GC-content dependent organizational tendency. To distinguish between these possibilities, we quantified correlations between GPSeq score and GC-content along chromosome arms after progressively removing bins near anchored landmarks of the genome (*CEN*, *TEL* & *rDNA*; **Supplementary Fig. 5a**). Removing the first ∼150 kb around these landmarks does not appreciably affect the correlations in either condition, indicating that the effect is not driven by telomeres becoming more central in A2Q (**Fig. 5d**). Interestingly, when we measured correlations between GC-content and GPSeq score for bins distant from either *TEL* or *CEN* (>200 kb), correlations steadily increased, reaching as high as SCC of 0.85 at the distance of 500 kb (**Fig. 5d**). While wt-Q similarly exhibits an initial increase in correlation, this increase plateaus (at SCC∼0.5) and subsequently declines sharply at ∼500 kb. (**Supplementary Fig. 5b**). By contrast, in cycling cells, we observed the same trend focusing only on *chrIVR*, the longest chromosome arm in the budding yeast genome (**Supplementary Fig. 5c-d**). These data reveal that DNA sequences far from anchor points in cycling cells follow the same behavior as the one observed in mammalian cells, where GC-rich sequences tend to be found toward the nuclear center^36,47^.

In order to test whether the GC-content of a locus can drive radial placement of DNA in the nucleus, we engineered yeast strains carrying a GC-poor insert (a 7.9 kb *Mycoplasma mycoides* fragment, 22% GC), positioned in a region with high GPSeq scores near the center of the long arm of chromosome II (*chrIIR*: *CEN* – 226 kb – *lys2*::insert – 354 kb – *TEL*; **Fig. 5e**). By placing a lac operator (lacO) array 5.9 kb from the insertion site and expressing LacI–GFP, we tracked the radial position of the locus relative to the nuclear envelope, marked with Nup49–mCherry (**Fig. 5e**). Introducing such GC-poor fragment resulted in a modest but statistically significant (p-val. = 0.0092) displacement of the locus toward the nuclear periphery (**Fig. 5f**), accompanied by an increase the proportion of cells where the locus is found at the nuclear envelope (p-val. = 0.0079, **Fig. 5g**). This provided initial evidence that local GC-content can exert a mild but detectable influence on the radial genome organization in yeast. The displacement however is milder than the effect of inducing heterochromatin spreading on a susceptible locus, shown previously^48^ (**Supplementary Fig. 5e-g**).

It has previously been shown that the recruitment of a locus to the nuclear periphery can lead to silencing of DNA loci, both in human and in budding yeast nuclei^8–10^. In yeast, this process was shown to occur in three steps^48^: (1) recruitment of a locus to the nuclear periphery where heterochromatin in concentrated; (2) this proximity to the peripheral heterochromatin and the associated silencing factors triggers the silencing of the locus; which in turn (3) stabilize the locus at the nuclear periphery^48,49^. We thus tested whether the peripheral movement of *LYS2* associated with the GC-poor insertion was sufficient to silence the silencing-susceptible cassette *E-ADE2-I* (**Fig. 5h**). Silencing of *ADE2* leads to the accumulation of a red pigment producing red or pink colonies depending on the strength of gene silencing, while colonies are white when *ADE2* is expressed^50,51^.

With only *E-ADE2-I* at *LYS2* the *ADE2* is not silenced in wild-type cycling cells, resulting in white colonies^48^ (**Fig. 5i**), in good agreement with the internal localization of this locus (as seen by microscopy and GPSeq) limiting contacts with the peripheral heterochromatin. However, with the GC-poor stretch of DNA integrated upstream of *E-ADE2-I*, we observed red colonies, indicating silencing of the *ADE2* gene (**Fig. 5i**). This finding strongly suggests that this GC-poor DNA fragment on its own is sufficient to position the locus peripherally and favor gene silencing (**Fig. 5j**).

## Discussion

In this work, we adapted Genomic Loci Positioning by Sequencing (GPSeq) assay to *Saccharomyces cerevisiae* to generate the first ever genome-wide high-resolution maps of radiality of the budding yeast genome. This approach enabled us to greatly enrich previous models of nuclear organization in yeast, which were mainly limited to the radial positioning of centromeres, telomeres, and the *rDNA*. This was possible thanks to the modifications to the GPSeq workflow we introduced, which made it allowed the study of cells with a cell wall, with nuclei of ∼ 1 um in radius, that were grown in suspension. We believe these modifications open up the possibility of adapting GPSeq to other model systems as well. We benchmarked GPSeq applied to yeast with previously published FROS-based radial measurements^33,34,42^ and observed a strong agreement between the two methods, validating our experimental approach. We note that GPSeq scores correlate better with datasets scoring the distance to the nuclear periphery^33^, than with those scoring for enrichment at the nuclear periphery^34^. This indicates that intermediate radial positions are accurately measure by GPSeq in the small budding yeast nucleus.

Centromeres and telomeres are known to localize predominantly at the nuclear periphery in the nucleus of cycling budding yeast cells, with the latter being most peripheral. As genomic distance from these landmarks increases, the distance to the nuclear periphery increases, consistent with the persistence length and compaction of the chromatin fiber in the budding yeast nucleus^37,40^. From GPSeq we learned how this specifically occurs over distances up to 100 kb from telomeres, and for up to 200 kb from centromeres. The distance from the centromeres matches the length scale of the polymer brush effect, previously shown to emanate from the centromere cluster^37^. Interestingly, for small chromosome arms, there is no inwards movement with increasing distance from the centromere, as telomere proximity seems dominant over the central-wards push associated with the centromere-associated polymer brush. Hence, small chromosomes are retained at the nuclear periphery. This is in line with previous models of the yeast 3D genome^52,53^ and measurements of telomere localization on small chromosomes (close to SPB)^54^. In summary, we find the radial placement of telomeres to dominate over centromere positioning and associated effects.

In conditions where telomeres are clustered in the center of the nucleus, the genome gets reorganized. Thanks to GPSeq we found that telomere clustering is accompanied by the centralization of a substantial portion of adjacent chromosome arms. This greatly affects short chromosome arms, as they are almost entirely dragged into the nuclear interior. At the same time, regions on the large chromosome arms that are distal from telomeres shift toward the nuclear periphery. We hypothesize that this configuration stems as a compensatory effect of hypercluster formation. The inwards movement of the telomeres into a ‘singular focus’, might impose a constraint that pushes chromosome arms outwards, through a polymer brush-like effect. This brush-like effect is more noticeable in A2Q than in wt-Q. Since additional rearrangements, such as long- and short-range DNA compaction, take place in the quiescent state^31,55^, they might account for these differences. Moreover, in the wt-Q cells, we also observed centromeres to lose their proximity to the nuclear periphery, which might release additional constraints on the chromosome arm. Indeed, centromere declustering has previously been reported in quiescent^31^ and stationary phase cells^56,57^. Based on all these observations, we propose two radial configurations to accommodate the genome in A2Q and wt-Q, as shown in Figure 3.

We next examined potential effects of transcription on genome organization and observed a mild enrichment of transcription and transcription-associated marks in the central portion of the budding yeast nucleus. This feature is observed across all conditions studied. However, within each condition, the relationship with transcription varied depending on chromosomal context. Mid-arm regions showed the clearest correlation between transcriptional activity and radial positioning, whereas this effect was weaker at telomere-proximal regions and absent near centromeres. In quiescence however, where centromeres are no longer at the very periphery and get declustered^31,56,57^, RNAPII distribution although sparse^27^ correlates better with genome radiality all along the genome than in wt. In wt-Q cells transcription is scant^27,28,30,31^, yet a fraction of RNAPII remains bound to the genome, ready to be activated at 30% of the promoters^27,58^. Here, we find that these regions are located relatively centrally in those cells. Such a spatial distribution might allow for the concentration of transcription-associated factors allowing for prompt reactivation of promoters upon return to growth.

Lastly, we found that GC-content as a genomic feature seems to correlate strongly with the variation in GPSeq scores within stretches of DNA that are not immediately adjacent to genomic anchor points. Often, genomic bins with relatively high GC content are located near chromosome ends (close to telomeres) or at the *rDNA* repeats that happen to be anchored at the nuclear membrane, which makes the interplay between the GC-content and radiality complex. We find that these anchored regions hide a tendency of chromatin to spatially organize with GC-content, which becomes increasingly apparent at greater genomic distances from the anchoring landmarks. Indeed, we observed a significantly higher genome-wide correlation between GC-content and the radial position of DNA loci upon detachment of the telomeres from the nuclear envelope in the A2Q cells. As a side note, the GC-content range across the genome is narrow^59^ (36–42% over 50 kb windows), indicating that relatively small changes in the GC-composition of a locus can have an observable effect.

We further show that introducing a 7.9 kb GC-poor stretch in the center of a chromosome arm is sufficient to displace the locus toward the nuclear periphery. Even more, integration of this GC-poor sequence upstream of a locus susceptible to heterochromatinization^48^, was sufficient to silence the locus. Notably, as the GC-poor DNA, we used a 7.9 kb fragment of *M. mycoides* DNA. Previously, Meneu *et al.* found that an entire *M. mycoides* chromosome (1220 kb long) integrated in the budding yeast nucleus forms a compact globule at the nuclear periphery^60^.

Our results extend these observations to a short sequence integrated far away from perinuclear anchors (centromeres or telomeres), while the full *M. mycoides* chromosome carried yeast centromeres and telomeres^60^. Our results show that the GC-poor sequence alone, is sufficient to bring a locus in proximity to the telomeric compartment, leading to its silencing. We propose several mechanisms to potentially explain the preferential localization of AT rich sequence at the nuclear peripheral. First, GC-poor sequences may not be optimal for the assembly of nucleosome in yeast leading to the formation of « perturbed chromatin » as shown for the *M. mycoides* chromosome in yeast^60^ and in *Neurospora Crassa* for native AT-rich DNA^61^. Perturbed chromatin could by itself trigger loci to the nuclear periphery as we previously showed^48^. Second, in human and budding yeast cells, AT-rich sequences, and repeats have been linked to increased replication stress^62,63^. In yeast both budding and fission yeast, certain types of replication stress can trigger the repositioning of affected loci to the nuclear periphery^64–66^. Lastly, AT base pairs are weaker and display reduced base stacking interactions, resulting in lower thermostability and increased bending rigidity of AT-rich DNA^67^. Because of this, AT-rich sequences behave as mechanically stiffer polymers, which polymer dynamics models suggest might bias them toward localization at the nuclear periphery^67,68^. In this framework, we note that the relatively small differences GC-content in budding yeast mirror broader GC-content variation in mammalian genomes, where ‘GC-poorer’, transcriptionally silent regions are preferentially positioned at the nuclear periphery^36,47,69^. We suggest that GC content may act as a conserved sequence-based cue for the spatial sequestration and silencing of DNA whose composition diverges from the mean of the genome, including potentially invasive sequences.

In conclusion, we adapted GPSeq to cycling and quiescent *S. cerevisiae* cells, whose nuclear radius is smaller than one micron, thereby extending its applicability to a broad range of additional organisms. We charted a radial map of the budding yeast genome and uncovered patterns that govern genome organization from *CEN* to *TEL*. In particular, our study unveils that GC content contributes to shaping the radial organization of the yeast genome. This property is shared with mammalian genomes^36,47,69^, suggesting a conserved mechanism.

## Materials and Methods

### Yeast strains used

All *S. cerevisiae* strains utilized in this section are haploid (mating type a) and share an isogenic background with W303 (**Supplementary Table 1**). Gene targeting for deletion and tagging was carried out following Longtine *et al.* (1998)^70^.

### Yeast culture and growth

Cells were initially inoculated in YPD (yeast-peptone-dextrose 2%) and cultured overnight. The following day, the cultures were diluted to 0.1 OD_600_/mL. The cells were grown at 30°C with shaking at 250 rpm in glass flasks, using 1/10 of the flask volume and aeration caps. Exponentially growing (cycling wt and cycling *SIR3-A2Q*) cells were harvested early in this timecourse at 1.5–3.5 OD_600_/mL. This stage is optimal for hypercluster formation in the *SIR3-A2Q* mutant. To induce quiescence through nutrient exhaustion (wt-Q, 6-7d-HD), cells were cultured for 6-7 days, until they reached stationary phase at ∼50 OD_600_/mL. For transcriptional silencing assays using the *ADE2*-based telomeric silencing reporter (*E-ADE2-I*), colonies were grown on YPD agar plates (yeast-peptone-dextrose 2%, agar 2%) at 30°C and subsequently incubated at 4 °C for 3–4 days to allow accumulation of the red pigment associated with *ade2* deficiency^50,51^.

### Crosslinking for GPSeq and enrichment of quiescent cells

To isolate the highly dense (HD) cells from *S. cerevisiae* stationary phase, samples taken at seven days were crosslinked and HD cells were enriched through gradient sorting. A total of 3.6×10^10^ cells (3000 OD_600_) were collected, resuspended in 1X PBS to a concentration of 1.8×10^8^ cells/mL (1 OD600/mL), and immediately crosslinked with 4% paraformaldehyde (16% stock solution, Pierce Thermo Scientific, 28906) for 15 minutes with shaking at room temperature. The reaction was quenched with 2.5 M glycine (final concentration 125 mM) for 5 minutes. Crosslinked cells were then pelleted by centrifugation at 800 g for 5 minutes, washed, and resuspended in 50 mM Tris pH 7.5 for sorting. The same day, cycling cells were crosslinked and quenched in the same manner as stationary phase cells, then immediately stored overnight in 1X PBS at 4°C.Following the method described by Allen et al. (2006), the stationary phase crosslinked cells were loaded onto a pre-established salt gradient. A solution of 150 mM NaCl in Percollwas prepared and distributed over 24 x 5 mL ultraclear Beckman Coulter tubes and centrifuged in a SW40Ti rotor at 19000g for 20 minutes. Subsequently, 140 OD_600_ units of crosslinked cells were loaded per gradient and centrifuged at 400g for one hour in swing buckets. The lower fraction, enriched in highly dense cells, was collected and washed twice with 1X PBS pH 7.6. Afterwards, the cells were stored overnight in 1X PBS at 4°C.

### Microscopy and image analysis

Images were acquired using a wide-field microscopy system controlled by MetaMorph software (Molecular Devices), built around an inverted Nikon TE2000 microscope with a 100X/1.4 NA immersion objective. The images were captured with a C-mos camera (ORCA-flash C11440, Hamamatsu, Japan) and illuminated by a Spectra X light engine lamp (Lumencor, Inc, Beaverton, OR, USA). This setup enables rapid acquisition of dual-color images when combined with a double filter. The displayed images are maximum intensity z-projections of z-stacks taken with a 200 nm step. For each experiment, all images were captured using the same acquisition parameters. Images were analyzed in Fiji^71^. When deconvolved, the Deconwolf software^72^ was used, with 25 iterations.

#### Radial nuclei segmentation and quantifications for GPSeq

To allow quantification of GPSeq digestion peripherality (YFISH), deconvolved images of DAPI and YFISH signal were used. The nuclear slice with the sharpest signal was taken, from which pixels were extracted using a cross-section (nuclear diameter) that did not include the rDNA. About 50 such cross-sections/diameters were taken per sample. We used a homemade algorithm to find the center and borders of the DAPI per nucleus. Then the border was taken as reference, giving two points per nucleus. On this, we overlaid YFISH signal and quantified peripherality of the restriction digestion.

#### Nuclei segmentation and quantifications for FROS

Whole nuclei were segmented and fluorescent locus positions determined using NuFoQal^73^, which derives the nuclear contour from the Nup49–mCherry signal and computes the distance of the *LacO*-bound LacI–GFP focus to this contour. This was done for three different conditions: for ‘no insert’ (*lys2::lacO)*, n=5, with rep. 1 = 541 cells, rep. 2 = 1394 cells, rep. 3 = 390 cells, rep. 4 = 238 cells, rep. 5 = 504 cells; for ‘GC-poor’ (*lys2::lacO M. mycoides)*, n=5, with rep. 1 = 534 cells, rep. 2 = 731 cells, rep. 3 = 500 cells, rep. 4 = 578 cells, rep. 5 = 627 cells; for ‘heterochromatinized’ (*lys2::lacO E-ADE2-I)*, n=3, with rep. 1 = 335 cells, rep. 2 = 501 cells, rep. 3 = 406 cells.

### GPSeq experimental procedures

#### Cell wall digestion and permeabilization

A single GPSeq digestion sample required a minimum of 80 OD_660_ (9.6×10^8^ cells) of (expo/cycling) cells or 160 OD_660_ (1.92×10^9^ cells) of quiescent cells. For cell wall removal, 20 OD_660_ cells were resuspended in 1.25 mL of 0.1M EDTA-KOH/EDTA-NaOH with 10 mM DTT and incubated at 30°C for 10 minutes. After centrifugation, cells were resuspended in 1.25 mL of YPD with 1.2M sorbitol and homogenized thoroughly. The cell wall lysing enzyme Zymolyase-100T (0.06 mg/mL, MP Biomedicals, 8320932) was added, and cells were incubated for 10 minutes for expo cells and 60 minutes for quiescent cells. For quiescent cells, Lyticase (0.06mg/mL; Sigma, L2524) was added at 30 minutes and incubation was prolonged for another 30 minutes. The reaction was stopped by adding 40 mL of YPD 1.2M sorbitol, followed by centrifugation. The cells were then washed twice with YPD 1.2M sorbitol and finally resuspended in 1X PBS.

#### Permeabilization and chromatin loosening

Permeabilization and chromatin loosening was executed in 50mL Falcon tubes for expo cells, while it was executed on coverslips for quiescent cells. Cells were incubated with 1X PBS with 0.5% Triton X-100 for 15 minutes at RT while shaking. After centrifugation, cells were resuspended in 1 mL of 1X PBS 20% glycerol and incubated at RT for at least 2 hours. Cells were flash-frozen in liquid nitrogen and thawed at RT, a process repeated four times. Following the freeze-thaw cycles, cells were washed with 1X PBS 0.05% Triton X-100, centrifuged, and then resuspended in Milli-Q water. Cells were treated with 0.1M HCl for 5 minutes. The reaction was stopped with an equal volume of 0.2M NaOH and then washed with 1X PBS with 0.05% Triton X-100 for another 5 minutes before being resuspended in 1X PBS.

#### Attachment of cells to coverslips

Coverslips were coated with Poly-L-Lysine (0.01-0.1%; Sigma, P8920). 50μL of cell suspension was added to individual coverslips. The cell suspension settled for 30 minutes on the coverslip, after which coverslips were spun down in six well plates at 850g. Afterwards, the remaining volume was washed away by 1X PBS and cells were kept in 1X PBS for several minutes.

#### DNA restriction digestion

Samples were washed with 2X SSC (saline-sodium citrate, Invitrogen, AM9770) and incubated in 2X SSC, 50mM Phosphate buffer, 50% Formamide solution at RT for 20 hours. Post-incubation, cells were washed with 2X SSC and 3.1 buffer (NEB, B7203S). A digestion mix was prepared with DpnII buffer, nuclease-free water (NFW) and different concentrations of DpnII enzyme (NEB, R0543M, **Supplementary Table 2**), pre-warmed at 37°C. Coverslips are incubated for in situ restriction digestion at 37°C for the varying times (Supplementary Table 2), followed by immediate transfer to ice-cold 1X PBS/50mM EDTA/0.01% Triton X-100 and washing multiple times on ice.

#### DNA Y-FISH

The following next steps were executed in a manner resembling what was described by Girelli et al. (2020). Briefly, the samples were dephosphorylated by incubating them in 400μl of 1X calf intestinal alkaline phosphatase buffer with 6μl of calf intestinal alkaline phosphatase (Promega, cat. no. M1821) for 2 hours at 37°C. Following this, we ligated YFISH adaptors (Supplementary methods Table 2) at a final concentration of 0.2μM in 300μL of 1X T4 DNA ligase buffer containing 36μL of T4 DNA ligase (Thermo Fisher Scientific, cat. no. EL0014) by incubating the samples for 18 hours at 16°C. The next day, we removed unligated adaptors by incubating the samples in 10mM Tris-HCl/1M NaCl/0.5% Triton X-100 pH 8, five times for 1 hour each at 37°C, while shaking. To prepare the hybridization mix, we diluted the labeled oligonucleotide to 200 nM in a hybridization buffer consisting of 2X SSC/25% formamide/10% dextran sulfate/1mg/ml E. coli tRNA/0.02% bovine serum albumin (BSA). We then placed the coverslips onto a piece of Parafilm, with cells facing a 300μl droplet of hybridization mix, and incubated the samples in a humidity chamber for 18 hours at 30°C. The following day, we washed the samples with 2X SSC 0.2% Tween-20 and incubated for seven minutes in 0.2X SSC 0.2% Tween-20 in a water bath at 45°C. The samples were washed once, and the seven minutes incubation was repeated. Afterwards, the samples were washed and incubated with 4X SSC 0.2% Tween-20. Finally, we incubated the samples in 2X SSC/0.1 ng/μl Hoechst 33342 (Thermo Fisher Scientific, cat. no. H3570) for 30 minutes at RT, rinsed them trice in 2X SSC, and mounted them with home-made mounting medium (2X SSC; 10mM Tris-HCl pH 7.5; 0.4% Glucose; 20mM Trolox, Sigma, 238813; 37μg/mL Glucose oxidase, Sigma, G2133; 1.2U/μL Catalase, Sigma, C3515) before imaging. All the samples were imaged as previously mentioned.

#### Library preparation

Library preparation was executed globally as in Girelli et al. (2020). The same procedure was used, as the one leading up to DNA-YFISH. We digested DNA, ligated the GPSeq adaptors^35^ (**Supplementary Table 3**), and washed unligated adaptors. We then scraped the cells off the coverslips and digested them in 110μl of 10mM Tris-HCl/100 mM NaCl/50 mM EDTA/1% SDS pH 8 containing 10μl of Proteinase K (NEB, cat. no. P8107S) for 18 hours at 56°C. The next day, we inactivated the enzyme by heating the mixture to 96°C for 10 minutes. We purified genomic DNA (gDNA) using phenol-chloroform extraction and precipitated the gDNA using glycogen (Sigma, cat. no. 10901393001) and sodium acetate, pH 5.5, in ice-cold ethanol (VWR, cat. no. 20821.310) for 18 hours at −80°C. The DNA pellets were resuspended in 100μl of TE buffer and sonicated using microTUBE AFA Fiber Pre-Slit Snap-Caps (Covaris, PN520045) in the Covaris E220 sonicator with the following settings: peak incident power 140W, duty factor 10%, 200 cycles per burst, 55s treatment time. We then concentrated the gDNA to a final volume of 8 μl in nuclease-free water using AMPure XP (Beckman Coulter, cat. no. A63881). Each sample was subjected to in vitro transcription separately with the MEGAscript T7 Transcription kit (Thermo Fisher Scientific, cat. no. AM1334-5), using 10 to 100 ng of gDNA (**Supplementary Table 3**) in a final volume of 20 μl, and incubating for 14 hours at 37 °C. After IVT, we added 1 μl of TURBO DNase (Invitrogen, AM2238) to each sample and incubated for 15 minutes at 37 °C. The RNA was then purified using Agencourt RNAClean XP beads (Beckman Coulter, cat. no. A63987). Finally, we prepared sequencing libraries with the TruSeq Small RNA Library Preparation kit (Illumina, cat. no. RS-200-0012), following the manufacturer’s instructions with some modifications. All libraries were sequenced on the NextSeq 500 system (Illumina) using the NextSeq 500/550 High Output v2 kit (75 cycles) (Illumina, cat. no. 20024906).

### Genome-wide data analysis

#### GPSeq preprocessing

GPSeq sequencing data was pre-processed using a yeast adapted version of the custom pipeline (gpseq-seq-gg^36^) featuring: quality control, read filtering based on the expected adaptor sequence, adaptor trimming, mapping, filtering of the mapping output, filtering of reads mapped away from restriction sites, and UMI-based read de-duplication. To consider multimapped reads, the sambamba function as in gpseq-seq-gg^36^, was altered, to not include mapping_quality in the filtering step. Two different reference genomes were used. For analyses related to chromosome landmarks (*CEN*, *TEL*, *rDNA*), genetic annotations (LTR retrotransposons, *LYS2* localization) and sequence features (GC content), a W303 reference genome was used (W303.asm01.HP0^41^). For comparisons of GPSeq scores with ChIP-ChIP and ChIP-seq datasets, the SacCer3 reference genome was used. Genome annotations were retrieved from associated GFF or GFF3 files. Outliers were not removed as in gpseq-seq-gg, but rather, bed files were saved and combined after deduplication, to be further processed using the R software environment (R version 4.2.1 (2022-06-23), https://www.rproject.org).

GPSeq score calculations were executed on a per cutsite basis, before binning at different scales. First, the proportion of reads (N_R_) allocated to a cutsite (cs) in a digestion (tD) was determined, P(cs, tD). Second, GPSeq score was calculated by dividing the proportion of reads allocated to a cs in the full digestion, P(cs, tFD), by the average proportion of reads found for that cutsite in the short digestions, μ(P(cs, tSD1-3)). See below.

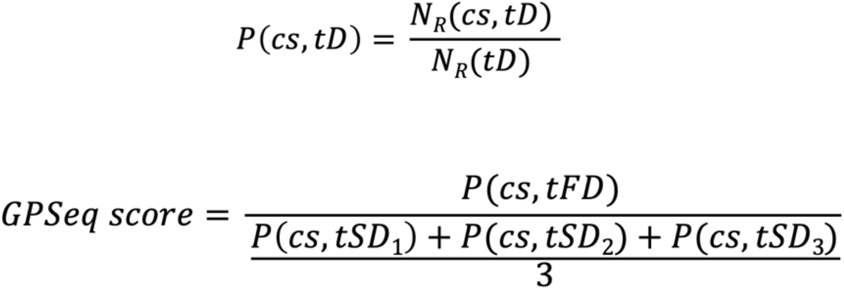

As an exception, for the second quiescent replicate, GPSeq scores were calculated using the full digestion from the first replicate together with the short digestions from the second replicate. This was necessary because the full digestion from the second replicate was of poor quality, showed weak correlation with the first replicate, and produced GPSeq scores that strongly diverged from those of the first replicate. Short digestions on the contrary behaved as expected.

GPSeq scores were transformed genome-wide, depending on the analysis in question: log_2_(GPSeq scores) and GPSeq scores were used to visualize the behaviour of GPSeq scores themselves. A z-score (GPSeq score – median(GPSeq score))/std(GPSeq score), is used to obeserve GPSeq score fluctuations relative to the median GPSeq score. In combination with z-scores, ranked GPSeq scores (percentiles) are also used to show positioning of regional GPSeq scores relative to the rest of the data. In addition, z-scores and percentiles were used to compare between conditions.

Computationally, in brief, bed files were concatenated in R and GPSeq scores were calculated, using the following packages: data.table^74^ and tidyverse^75^. After GPSeq score calculation lower and upper outliers were removed using the outliers package^76^. Genome-wide binning (sliding, overlapping & non-overlapping) was executed using homemade binning algorithms, relying on the GenomicRanges^77^, IRanges^77^ and GenomeInfoDb^78^ packages for genomic interval representations and overlap calculations.

### Plotting and statistical analyses

Plotting and statistical analyses were performed using the R software environment. The packages ggplot2^79^, ggsignif^80^ and pheatmap^81^ were used, alongside base R functions. For statistical analysis, the specific method used depended on the application. Non-parametric wilcoxon tests were used to compute significance between two groups. When multiple groups were compared (e.g. FROS-analyses; GPSeq *TEL* and *CEN* comparisons) either linear and generalized linear mixed-effects models were fitted using the lmer and glmer functions from the lme4 package^82^ to account for fixed and random effects.

### Micro-C contact data

Reads were aligned and the contact data processed using Hicstuff^83^ (version 3.2.4), available on Github (https://github.com/koszullab/hicstuff). Briefly, pairs of reads were aligned iteratively and independently using Bowtie2^84^ in its most sensitive mode against the *S. cerevisiae* reference genome (W303.asm01.HP0^41^). Each uniquely mapped read was assigned to a restriction fragment. Quantification of pairwise contacts between restriction fragments was performed with default parameters: uncuts, loops and circularization events were filtered as described previously^36^. PCR duplicates (defined as paired reads mapping at exactly the same position) were discarded. Contact maps were binned at the resolution of 2 kb and normalized using balance functions of Cooler. To compute and visualise the agglomerated plots, we used the algorithm Chromosight^85^ in quantify mode (version 1.6.3, available on Github (https://github.com/koszullab/chromosight) with the following options (--win-size=401 --perc-undetected=100 --perc-zero=100).

### Comparison with other datasets

All data sets were lifted over to SacCer3 when required. Histone marks data was obtained from Weiner *et al.* (2015) and processed as described in Hocher *et al.* (2018). Sir3 binding in cycling cells and quiescent cells was obtained from Hocher *et al.* (2018) and Guidi *et al.* (2015) respectively. RNA-seq reads from Hocher *et al.* (2018) and Baquero Pérez *et al.* (2025) were trimmed (using TrimGalore^86^) and mapped to the Saccer3 assembly with Hisat^87^ 2.2.1 (HISAT2 –max-intronlen 500). CPM normalized coverage files without duplicates were generated with bamCoverage from deepTools^88^ (3.5.0). RNA polymerase II datasets from Baquero Pérez *et al.* (2025), and ATAC-seq data from Hsu C. *et al.* Gene Expression Omnibus sample GSM5172250. NCBI GEO (2021). Comparisons between datasets were performed exclusively with sliding windows of 45 kb (5 kb step).

## Acknowledgements

We acknowledge the members of the Taddei laboratory, UMR3664 and the MSCA ITN Cell2Cell for helpful discussions. We thank Nicola Crosetto for advice and comments on the manuscript. The A.T. team was financially supported by funding from Agence Nationale pour la Recherche DeSynLE (ANR-22-CE120013-01) and Labex DEEP (ANR-11-LABEX-0044 DEEP and ANR-10IDEX-0001-02 PSL). The authors greatly acknowledge the PICT-IBiSA@Pasteur Imaging Facility of the Institut Curie, a member of the France Bioimaging National Infrastructure (ANR-24-INBS-0005 FBI BIOGEN). G.L. and W.H.Y. were supported by a PhD fellowship from the European Union’s Horizon 2020 Research and Innovation programme under the Marie Skłodowska-Curie Actions Innovative Training Network (MSCA ITN ‘Cell2Cell’ – grant no. 860675). G.L. and M.B.P. were also supported by Labex DEEP (ANR-11-LABEX-0044 DEEP). In addition, this work was funded through research grants from the Swedish Research Council (grant. no. 2020-02657), Karolinska Institutet (KI Consolidator Grants 2020), and the European Union (ERC, RADIALIS, GA n. 101088408) and intramural funding from Human Technopole to M.B. Views and opinions expressed are those of the authors only and do not necessarily reflect those of the European Union or the European Research Council Executive Agency. Neither the European Union nor the granting authority can be held responsible for them.

## Author Contributions Statement

Conceptualization: A.T., M.B., G.L. Data curation: G.L., W.H.Y., M.B.P. Sample preparation: G.L. Sequencing data acquisition: W.H.Y. FISH validation and analysis: G.L. FROS and ESA experiments and analysis: G.L. Formal analysis: G.L., M.B.P., A.C. Resources and Funding acquisition: A.T., M.B. Investigation: G.L., W.H.Y., M.B.P., A.C., A.T., M.B. Project administration: G.L., A.T., M.B. Supervision: A.T., M.B. Visualization: G.L., M.B.P., A.C. Writing - original draft: G.L., A.T., M.B. Writing - review and editing: all authors.

