## supplementary Information for "A radial map of the budding yeast genome reveals novel organizational principles"

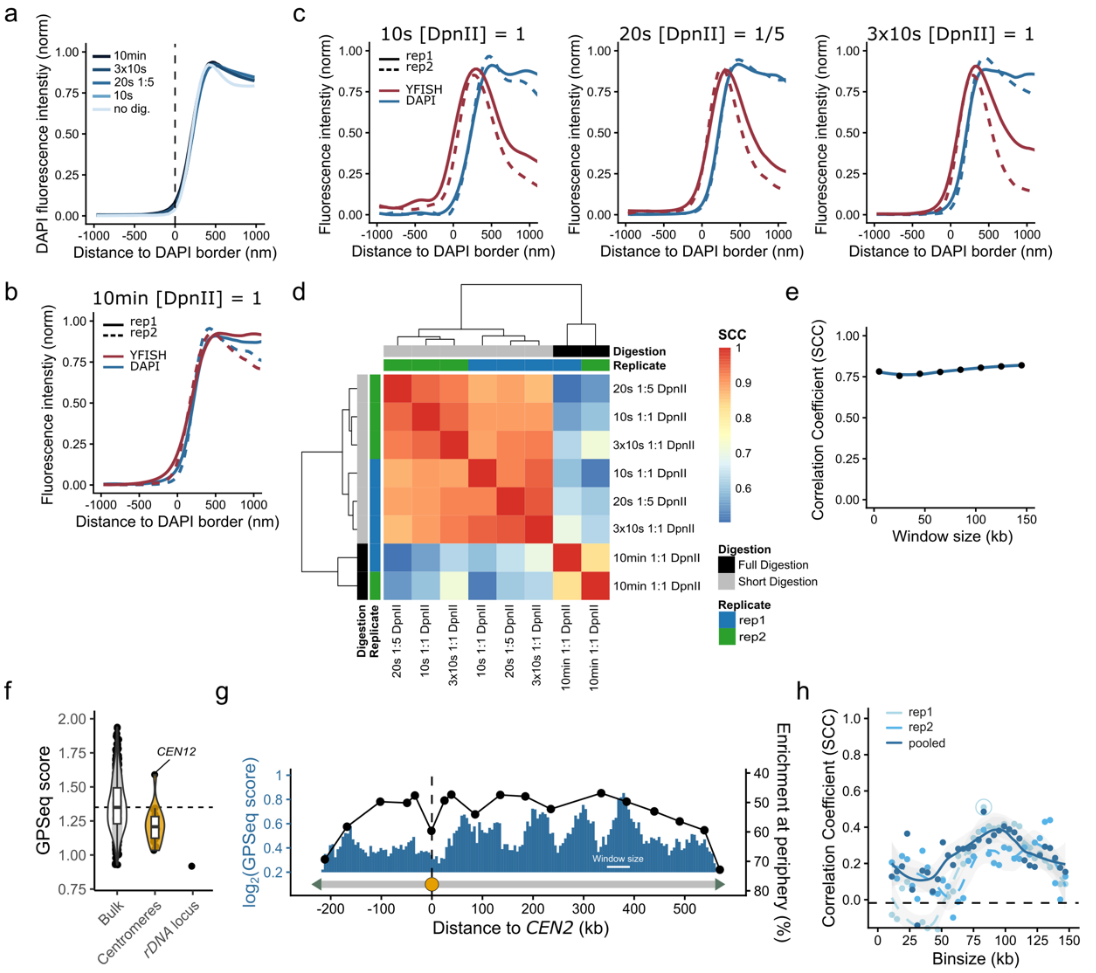


**Supplementary Figure 1: GPSeq optimization in *S. cerevisiae*. a**: Identification of the nuclear border, which was segmented using DAPI signals. **b**: Normalized YFISH signal intensity relative to DAPI signal for a full digestion of 10 min, for two biological replicates. **c**: Normalized YFISH signal intensity relative to DAPI signal for different short digestions of 10s, 20s (diluted 1:5) and 3x10s, for two biological replicates. **d**: Hierarchical clustering of Spearman Correlation Coefficient (SCC) matrix, showing correlations between normalized read counts for the different digestions. **e**: Progression of SCC between the two biological replicates across different window sizes. **f**: GPSeq scores for the different centromeres and *rDNA* locus compared to the bulk at 20 kb resolution, *CEN12* is highlighted. For the *rDNA* locus, multimapping reads were allowed. **g**: Chromosome profile of *chrII*, combining $\log_2$(GPSeq scores) with the point-median data of Dultz *et al.* (2016) superimposed. GPSeq score is plotted in 45 kb windows, sliding in 5 kb steps. **h**: SCC between GPSeq scores and loci probed through FROS-imaging for chrII by Dultz *et al.* (2016), for increasing binsizes. Size of circles surrounding each point represented the $-\log_{10}$(p-value) associated with the correlation, only when p.value < 0.05.

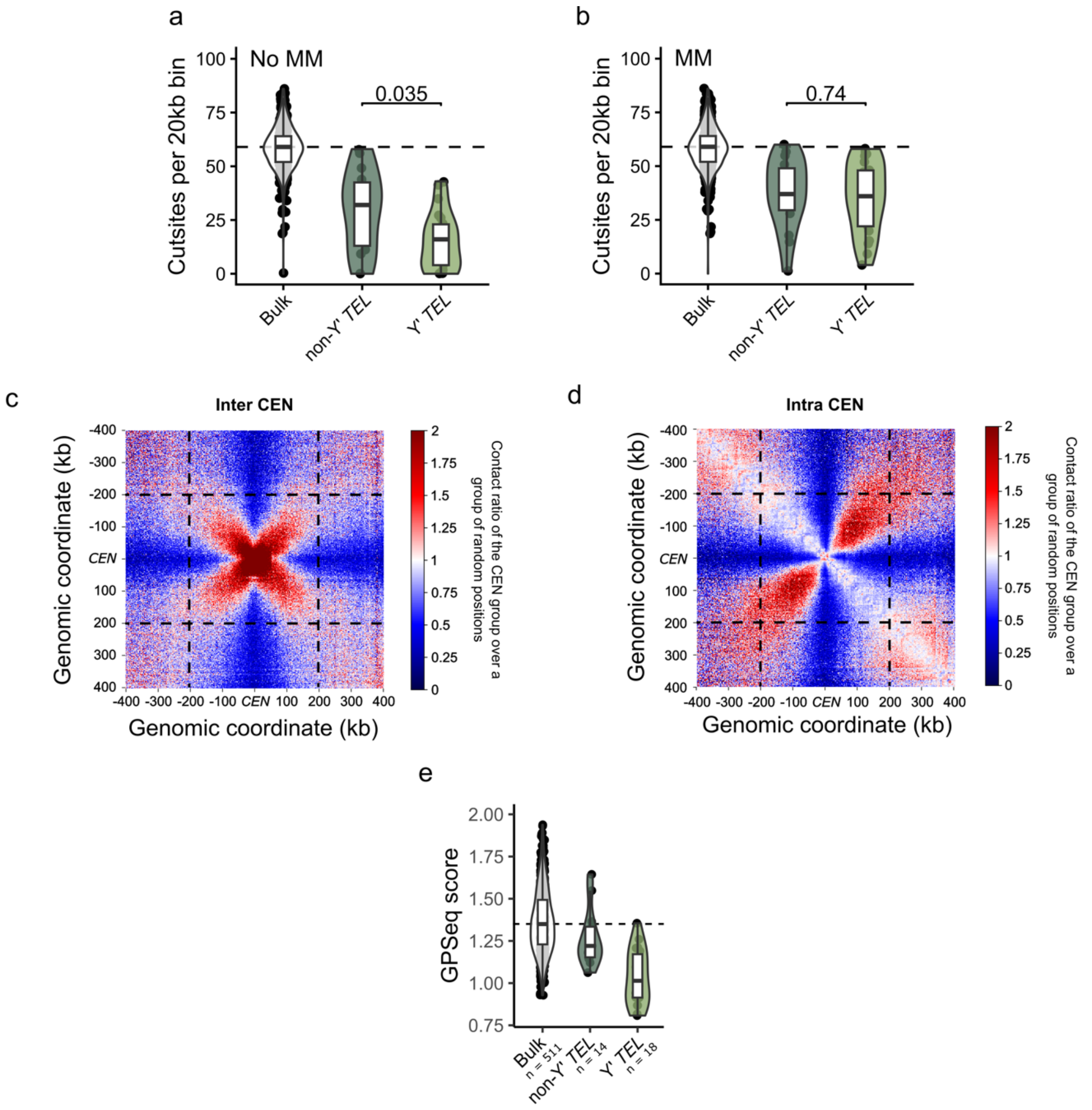


**Supplementary Figure 2: The effect of centromeres and telomeres on genome organization. a**: Amount of cutsites, containing at least one read, found in 20 kb bins when multimapping reads are not allowed. **b**: Amount of cutsites, containing at least one read, found in 20 kb bins, allowing multimapping reads for the telomeric most 13 kb. **c**: Agglomerated plot of the 120 contact sub-matrices between centromeric regions (*CEN*) from different chromsomes for asynchronous cells in log phase (MicroC data from Swygert *et al.* (2021)2) detrended by expected values. **d**: Agglomerated plot of the 16 contact sub-matrices at the centromeric regions (*CEN*) for asynchronous cells in log phase (MicroC data from Swygert *et al.* (2021)) detrended by expected values. **e**: GPSeq scores of bins associated with telomeres either containing (Y' *TEL*) or not containing (non-Y' *TEL*) a repetitive Y' element. Bins are compared to the bulk at 20 kb resolution.

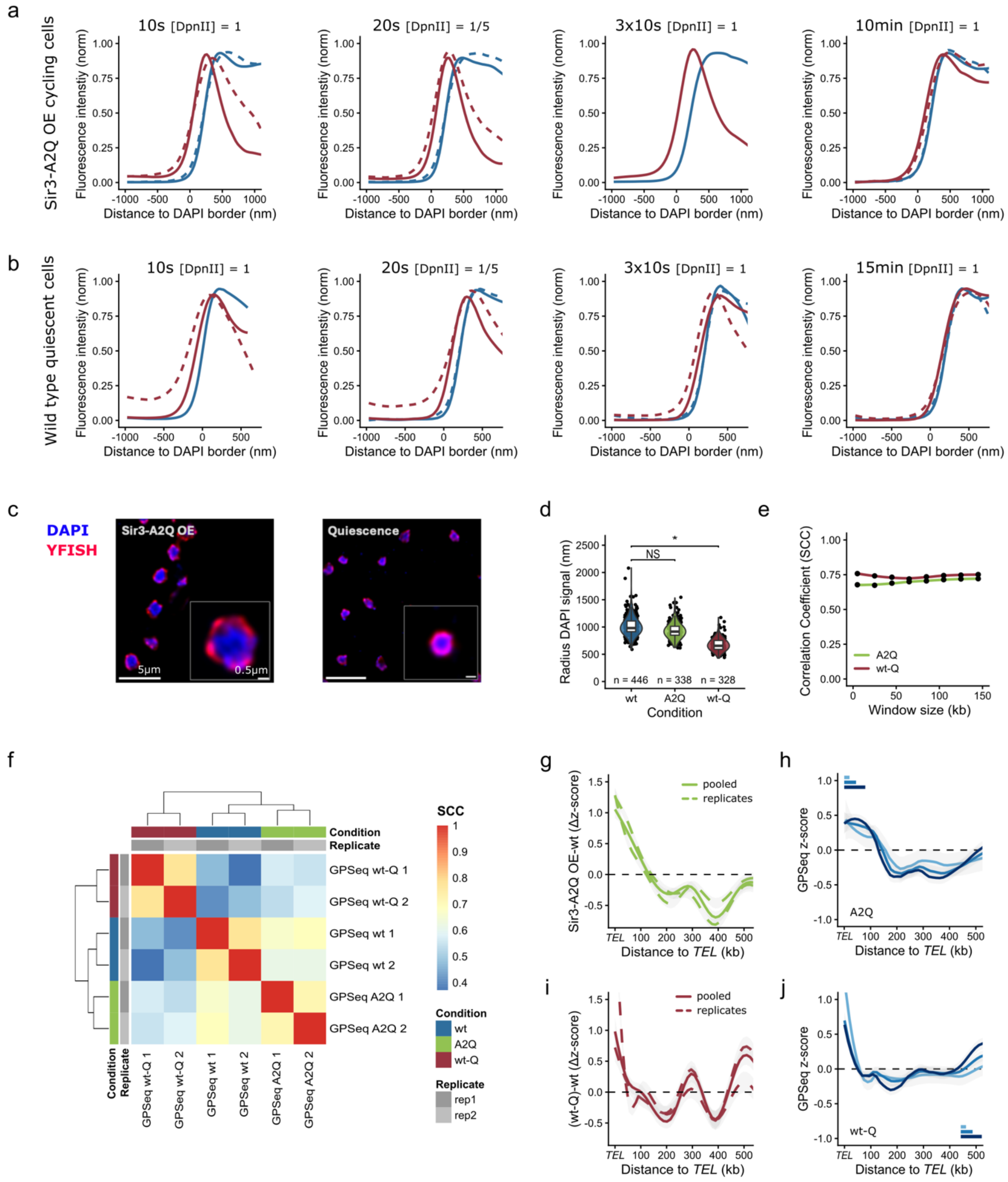

a
Sir3-A2Q OE cycling cells
10s [DpnII] = 1
20s [DpnII] = 1/5
3x10s [DpnII] = 1
10min [DpnII] = 1
Fluorescence intenstiy (norm)
Distance to DAPI border (nm)
b
Wild type quiescent cells
10s [DpnII] = 1
20s [DpnII] = 1/5
3x10s [DpnII] = 1
15min [DpnII] = 1
c
DAPI
YFISH
Sir3-A2Q OE
Quiescence
5μm
0.5μm
d
Radius DAPI signal (nm)
NS
*
n = 446
n = 338
n = 328
wt
A2Q
wt-Q
Condition
e
Correlation Coefficient (SCC)
A2Q
wt-Q
Window size (kb)
f
SCC
Condition
Replicate
GPSeq wt-Q 1
GPSeq wt-Q 2
GPSeq wt 1
GPSeq wt 2
GPSeq A2Q 1
GPSeq A2Q 2
wt
A2Q
wt-Q
rep1
rep2
g
Sir3-A2Q OE-wt (Δz-score)
pooled
replicates
Distance to TEL (kb)
h
GPSeq z-score
A2Q
i
(wt-Q)-wt (Δz-score)
j
wt-Q

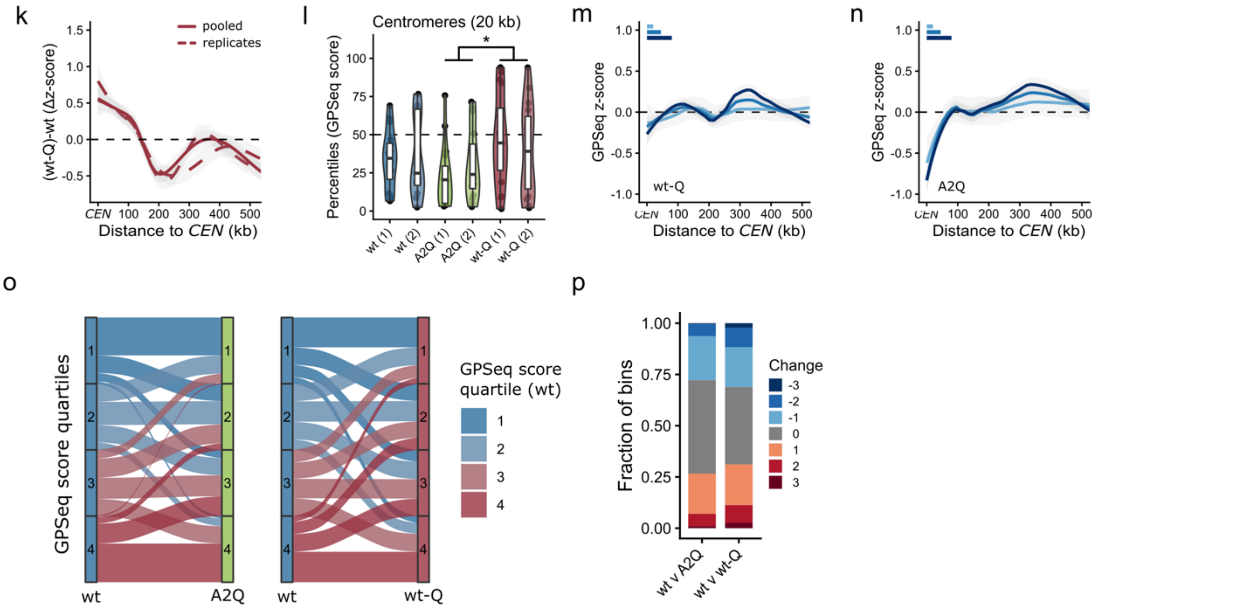


**Supplementary Figure 3: GPSeq in A2Q and Q: Global changes during telomere centralization.** **a**: Normalized YFISH signal intensity relative to DAPI signal for the A2Q condition, visualized for two biological replicates. **b**: Normalized YFISH signal intensity relative to DAPI signal for the wt-Q condition, visualized for two biological replicates. **c:** Representative images for short digestions for A2Q and Q (20s [DpnII] = 1/5). **d:** Distribution of DAPI signal Radii as indication for nuclear size, measured from border-to-border; NS: p.val > 0.05, *: p.val < 0.05. **e**: SCCs between different biological replicates for the different conditions (A2Q:green, wt-Q:red). **f**: Hierarchical clustering of SCC matrix for calculated GPSeq scores, showing how biological replicates cluster together within conditions. **g**: Average difference in GPSeq z-score between A2Q and wt visualized for the first 500 kb adjacent to telomeres. GPSeq scores are shown for 80 kb windows, sliding in 5 kb steps. **h**: Average GPSeq z-scores extending from the telomere for A2Q. Line color in represents different window sizes (scales relative to x-axis). Smallest binsize shows non-overlapping (20 kb) windows, while the other two represent overlapping sliding windows, respectively 45 kb and 80 kb, both with a 5 kb step. **i**: Average difference in GPSeq z-score between wt-Q and wt visualized for the first 500 kb adjacent to telomeres. GPSeq scores are shown from 80 kb windows, sliding in 5 kb steps. **j**: Same as in **h**, but for wt-Q. **k**: Average difference in GPSeq z-score between wt-Q and wt visualized for the first 500 kb adjacent to a centromere. GPSeq scores are shown for 80 kb windows, sliding in 5 kb steps. **l**: GPSeq score percentiles for centromeric reads (20 kb bins) showing replicates for each condition. Different conditions are colored (wt:blue, A2Q:green, wt-Q:red). Pairwise significance was assessed from estimated marginal means derived from a generalized linear mixed-effects model, with biological replicate included as a random effect: *: p-value < 0.05. **m**: Same as in **j** but now showing average GPSeq z-scores extending from the centromere. **n**: Same as in **m**, but for A2Q. **o**: Alluvial diagram showing radial movement of bins between conditions, numbered rows represent quartiles of GPSeq score (1: most peripheral, 4: most central). 80 kb sliding windows were used with a 5 kb step. **p**: Quantification associated with **o**, showing proportion of all bins associated with movement into a more peripheral or central quartile. 0: stays in same quartile, 1 to 3: movement of bin to more central quartile, -1 to -3: movement of bin to more peripheral quartile.

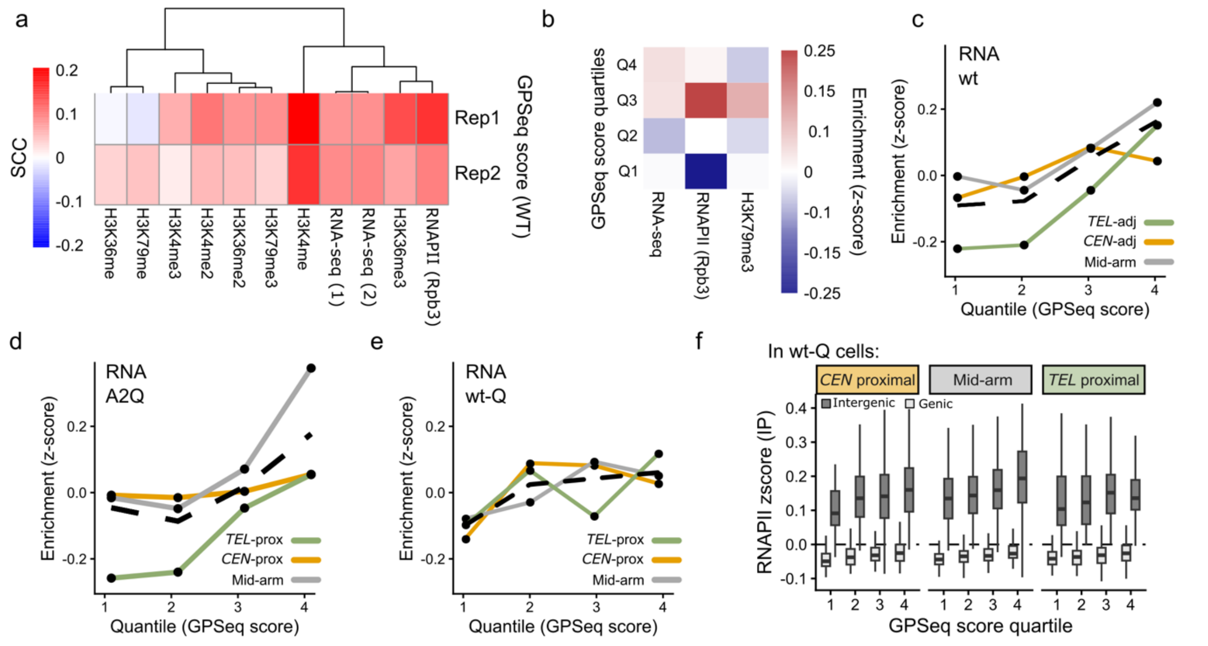


**Supplementary Figure 4: Enrichment of active transcription in the center of the nucleus. a**: SCCs between GPSeq score and features associated with transcription. Correlations were assessed at sliding window sizes of 45 kb, with 5 kb steps. Histone modification data was used from Weiner *et al.* (2015)[3], RNA-seq data from Hocher *et al.* (2018)[4] and RNAPII data from Baquero Pérez *et al.* (2025)[5]. **b**: Average feature z-score per GPSeq score quartile for wt. *TEL*-proximal 150 kb is excluded. Calculated from 45 kb sliding windows with 5 kb steps). H3K79me3 data was used from Weiner *et al.* (2015), RNA-seq data from Hocher *et al.* (2018) and RNAPII data from Baquero Pérez *et al.* (2025). **c-e**: Average RNA-seq counts z-score over different GPSeq score quartiles for wt, A2Q and wt-A2Q. Calculated from 45 kb sliding windows (5 kb step). Bins are categorized by their proximity to *CEN* (634 bins)/ *TEL* (636 bins)/ mid-arm (1025 bins). Dashed line represents the genome-wide average. RNA-seq data for wt and A2Q was used from Hocher *et al.* (2018) and RNA-seq data in wt-Q from Baquero Pérez *et al.* (2025). **f**: Boxplots showing differences in RNAPII (Rpb3) IP counts z-score for both intergenic and genic sections of the genome in wt-Q cells. Shown for the different sections of the chromosome arm. RNAPII data in wt-Q was used from Baquero Pérez *et al.* (2025).

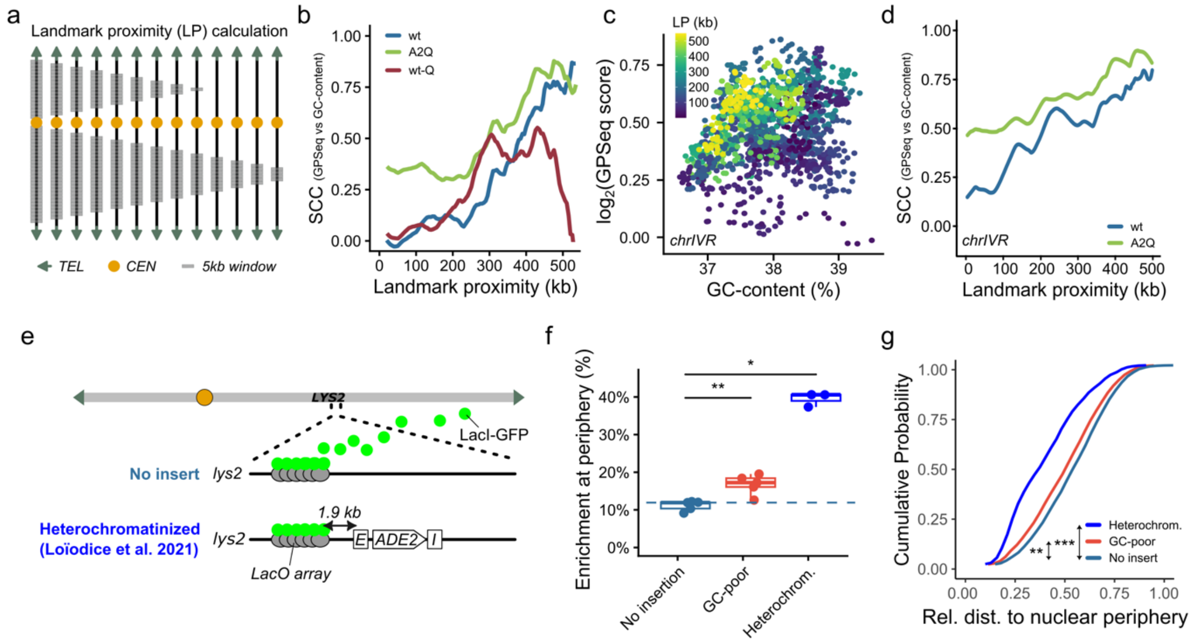


**Supplementary Figure 5: GC-poor DNA facilitates modest movement of chromatin towards the nuclear periphery. a**: Scheme accompanying Fig. 5D, showing sequential removal of bins near chromosome landmarks *CEN* and *TEL*. **b**: Correlation between GC-content and GPSeq score, shown while sequentially removing bins nearest to chromosome landmarks (*CEN*, *TEL* and *rDNA*). SCCs are shown for 45 kb sliding windows (step: 5 kb). **c**: Scatterplot showing $\log_2$(GPSeq score) versus GC content for bins of *chrIVR* (45 kb sliding windows, step: 5 kb). Color scale shows landmark proximity (LP). **d**: SCCs between GC-content and GPSeq scores shown for *chrIVR* while sequentially removing bins nearest to chromosome landmarks (*CEN* and *TEL*). Correlation shown for 45 kb sliding windows (step: 5 kb). **e**: Schematic representation of *chrII* with *LYS2* locus, showing integration site at *lys2* used in Loïodice *et al.* (2021), for the FROS assay. **f**: Proportion of *lys2::lacO* (n=5), *lys2::lacO M. mycoides* (7.9 kb, n = 5) and *lys2::lacO E-ADE2-I* (yAT2059, from Loïodice *et al*. (2021); n =3) loci found at the nuclear periphery. The nuclear periphery is marked as the peripheral most third of the nuclear area. Significance is shown for the results of a Wilcoxon rank-sum test: *: p-value < 0.05; **: p-value < 0.01. **g**: Cumulative distributions of relative distance to the nuclear periphery for *lys2::lacO* (n=5), *lys2::lacO M. mycoides* (7.9 kb, n = 5) and *lys2::lacO E-ADE2-I* (yAT2059, from Loïodice *et al*. (2021); n =3) loci. Pairwise significance (Bonferroni post-hoc comparison) was assessed from estimated marginal means derived from a generalized linear mixed-effects model, with biological replicates and experiment included as a random effect: **: p-value < 0.01, ***: p-value < 0.001 .

**Supplementary Table 1: Strains used in this work.** FROS: Fluorescent Repressor-Operator System, ESA: Ectopic Silencing Assay

| Publication | Experiment | Strain | Genotype |
|---|---|---|---|
| | / | **W303** | *ade2-1 can1-100 his3-11,15 ura3-1 leu2-3,112 rad5- trp1-1* |
| **Loïodice *et al.* (2021)**[6] | **GPSeq** | yAT126 | *ade2-1::ADE2* |
| **Ruault *et al.* (2025)**[7] | **GPSeq** | yAT2487 | *ade2-1::ADE2 hml∆::HPH rap1::GFP-RAP1(LEU2)* |
| **Ruault *et al.* (2021)**[8] | **GPSeq** | yAT2555 | *Ade2-1::ADE2 hml∆::HPH rap1::RAP1-GFP(LEU2) sir3::GPDp-Sir3-A2Q(NAT)* |
| **Loïodice *et al.* (2021)** | **FROS/SA** | yAT2059 | *lys2::E-ADE2-I lacOfx 1 array (TRP1) nup49::mCherry-NUP49 (URA3) leu2:: LacIR-GFP (LEU2)* |
| **This work** | **FROS** | yAT4954 | *ade2-1::ADE2 Nup49::mCherry-NUP49-URA3 leu2::Hisp-GFP-LacIR (leu2::KanMX3) lys2:: lacOfx (TRP1)* |
| **This work** | **FROS** | yAT4955 | *ade2-1::ADE2 Nup49::mCherry-NUP49-URA3 leu2::Hisp-GFP-LacIR (leu2::KanMX3) lys2:: lacOfx (TRP1) lys2::M.Mycoides(7.9Kb)* |
| **Loïodice *et al.* (2021)** | **ESA** | yAT2956 | *lys2:: E-ADE2-I* |
| **This work** | **ESA** | yAT4957 | *lys2:: E-ADE2-I lys2::M.mycoides (7.9Kb)* |
| **This work** | **ESA** | yAT4958 | *lys2:: E-ADE2-I lys2::M.mycoides (7.9Kb)* |

**Supplementary Table 2: Different restriction digestions executed**

| Digestion type | DpnII concentration | DpnII digestion time |
|---|---|---|
| Short digestion | 0.5 U/µL | 10 seconds |
| Short digestion | 0.1 U/µL | 20 seconds |
| Short digestion | 0.5 U/µL | 3*10 seconds (each time blocked with ice cold 1X PBS/50mM EDTA/0.01% Triton X-100, then washed with 2X SCC and 1X DpnII buffer) |
| Full digestion | 0.5 U/µL | 10 minutes (15 minutes for quiescence) |

**Supplementary Table 3:** List of oligos to design YFISH and GPSeq sequencing adapters. All oligonucleotides were purified using desalting, except for C2_Atto647N, that was purified using HPLC. Barcodes for sequencing oligos are in green.

**Oligos for YFISH**

| **Oligo name** | **Sequence (5' → 3')** |
|---|---|
| YFISH_DpnII_C2_Sense | /5PHOS/GATCGTCTATCTAGGCTAGTTCATCAGCTACGTTACGGAGTTCGGGGAATCTTAAACCCAACTTGTTTGAATCTTAAACCCAACTTGT |
| YFISH_DpnII_C2_Antisense | GAATCTTAAACCCAACTTGTTTGAATCTTAAACCCAACTTGTCCCGAACTCCGTAACGTAGCTGATGAACTAGCCTAGATAGAC |
| C2_Atto647N | ACAAGTTGGGTTTAAGATTC/3ATTO647NN/ |

**Oligos for GPSeq**

| **Oligo name** | **Sequence (5' → 3')** |
|---|---|
| GPSeq_DpnII_1_Sense | /5PHOS/GATCGCGTGATGNNNNNNNNGATCGTCGGACTGTAGAACTCTGAACCCCTATAGTGAGTCGTATTACCGGCCTCAATCGAA |
| GPSeq_DpnII_1_Antisense | CGATTGAGGCCGGTAATACGACTCACTATAGGGGTTCAGAGTTCTACAGTCCGACGATCNNNNNNNN**CATCACGC** |
| GPSeq_DpnII_2_Sense | /5PHOS/GATCGCGACGACNNNNNNNNGATCGTCGGACTGTAGAACTCTGAACCCCTATAGTGAGTCGTATTACCGGCCTCAATCGAA |
| GPSeq_DpnII_2_Antisense | CGATTGAGGCCGGTAATACGACTCACTATAGGGGTTCAGAGTTCTACAGTCCGACGATCNNNNNNNN**GTCGTCGC** |
| GPSeq_DpnII_3_Sense | /5PHOS/GATCGCGGTCGTNNNNNNNNGATCGTCGGACTGTAGAACTCTGAACCCCTATAGTGAGTCGTATTACCGGCCTCAATCGAA |
| GPSeq_DpnII_3_Antisense | CGATTGAGGCCGGTAATACGACTCACTATAGGGGTTCAGAGTTCTACAGTCCGACGATCNNNNNNNN**ACGACCGC** |
| GPSeq_DpnII_4_Sense | /5PHOS/GATCGCGCATCANNNNNNNNGATCGTCGGACTGTAGAACTCTGAACCCCTATAGTGAGTCGTATTACCGGCCTCAATCGAA |
| GPSeq_DpnII_4_Antisense | CGATTGAGGCCGGTAATACGACTCACTATAGGGGTTCAGAGTTCTACAGTCCGACGATCNNNNNNNN**TGATGCGC** |
| GPSeq_DpnII_5_Sense | /5PHOS/GATCGATTGATGNNNNNNNNGATCGTCGGACTGTAGAACTCTGAACCCCTATAGTGAGTCGTATTACCGGCCTCAATCGAA |
| GPSeq_DpnII_5_Antisense | CGATTGAGGCCGGTAATACGACTCACTATAGGGGTTCAGAGTTCTACAGTCCGACGATCNNNNNNNN**CATCAATC** |
| GPSeq_DpnII_6_Sense | /5PHOS/GATCGATTGATGNNNNNNNNGATCGTCGGACTGTAGAACTCTGAACCCCTATAGTGAGTCGTATTACCGGCCTCAATCGAA |
| GPSeq_DpnII_6_Antisense | CGATTGAGGCCGGTAATACGACTCACTATAGGGGTTCAGAGTTCTACAGTCCGACGATCNNNNNNNN**GTCGTATC** |

**Supplementary Table 4:** Input DNA for *in vitro* transcription (IVT), for all conditions

| Strain | Condition | Digestion | Input DNA (ng/μL) |
|---|---|---|---|
| **yAT126** | Wild type (rep1) | 10s, 0.5 U/μL | 2.38 |
| **yAT126** | Wild type (rep1) | 20s, 0.1 U/μL | 2.10 |
| **yAT126** | Wild type (rep1) | 3*10s, 0.5 U/μL | 1.25 |
| **yAT126** | Wild type (rep1) | 10min, 0.5 U/μL | 0.76 |
| **yAT126** | Wild type (rep2) | 10s, 0.5 U/μL | 3.14 |
| **yAT126** | Wild type (rep2) | 20s, 0.1 U/μL | 1.73 |
| **yAT126** | Wild type (rep2) | 3*10s, 0.5 U/μL | 1.25 |
| **yAT126** | Wild type (rep2) | 10min, 0.5 U/μL | 0.496 |
| **yAT2555** | Sir3-A2Q OE (rep1) | 10s, 0.5 U/μL | 1.55 |
| **yAT2555** | Sir3-A2Q OE (rep1) | 20s, 0.1 U/μL | 1.22 |
| **yAT2555** | Sir3-A2Q OE (rep1) | 3*10s, 0.5 U/μL | 1.36 |
| **yAT2555** | Sir3-A2Q OE (rep1) | 10min, 0.5 U/μL | 1.14 |
| **yAT2487** | Q (7d-HD, rep1) | 10s, 0.5 U/μL | 4.34 |
| **yAT2487** | Q (7d-HD, rep1) | 20s, 0.1 U/μL | 4.32 |
| **yAT2487** | Q (7d-HD, rep1) | 3*10s, 0.5 U/μL | 4.86 |
| **yAT2487** | Q (7d-HD, rep1) | 15min, 0.5 U/μL | 1.16 |
| **yAT2555** | Sir3-A2Q OE (rep2) | 20s, 0.1 U/μL | 1.95 |
| **yAT2555** | Sir3-A2Q OE (rep2) | 3*10s, 0.5 U/μL | 2.31 |
| **yAT2555** | Sir3-A2Q OE (rep2) | 10min, 0.5 U/μL | 2.06 |
| **yAT2487** | Q (7d-HD, rep2) | 20s, 0.1 U/μL | 2.67 |
| **yAT2487** | Q (7d-HD, rep2) | 3*10s, 0.5 U/μL | 2.07 |
| **yAT2487** | Q (7d-HD, rep2) | 15min, 0.5 U/μL | 1.50 |
